# Statistical Curve Models For Inferring 3D Chromatin Architecture

**DOI:** 10.1101/2022.02.19.481149

**Authors:** Elena Tuzhilina, Trevor Hastie, Mark Segal

## Abstract

Reconstructing three dimensional (3D) chromatin structure from conformation capture assays (such as Hi-C) is a critical task in computational biology, since chromatin spatial architecture plays a vital role in numerous cellular processes and direct imaging is challenging. We previously introduced Poisson metric scaling (PoisMS), a technique that models chromatin by a smooth curve, which yielded promising results. In this paper, we advance several ways for improving PoisMS. In particular, we address initialization issues by using a smoothing spline basis. The resulting SPoisMS method produces a sequence of reconstructions re-using previous solutions as warm starts. Importantly, this approach permits smoothing degree to be determined via cross-validation which was problematic using our prior B-spline basis. In addition, motivated by the sparsity of Hi-C contact data, especially when obtained from single-cell assays, we appreciably extend the class of distributions used to model contact counts. We build a general distribution-based metric scaling (DBMS) framework, from which we develop zero-inflated and Hurdle Poisson models as well as negative binomial applications. Illustrative applications make recourse to bulk Hi-C data from IMR90 cells and single-cell Hi-C data from mouse embryonic stem cells.

## 1. Introduction

The task of reconstructing the three-dimensional (3D) configuration of chromatin (from a single chromosome) within the eukaryotic nucleus from pairwise contact assays, notably Hi-C (Lieberman-Aiden et al., 2009; Duan et al., 2010; Rao et al., 2014), is motivated by (at least) three considerations. First, such architecture is critical to an array of cellular processes, particularly transcription, but even memory formation (Marco et al., 2020). Second, armed with such an inferred configuration, we can superpose genomic attributes, enabling biological insights not accessible from the primary Hi-C contact matrix readout. Examples here include gene expression gradients and co-localization of virulence genes in the malaria parasite (Ay et al., 2014), the impact of spatial organization on double strand break repair (Lee et al., 2016), and elucidation of ‘3D hotspots’ corresponding to (say) overlaid ChIP-Seq transcription factor extremes which can reveal novel regulatory interactions (Capurso, Bengtsson and Segal, 2016). Third, despite notable gains in imaging methodologies (Payne et al., 2021), such direct access to structure is yet to enjoy the resolution and uptake conferred by Hi-C assays.

This set of factors has led to a wealth of 3D reconstruction algorithms: a recent review (Oluwadare, Highsmith and Cheng, 2019) identified over 30 methods and there have numerous additions in subsequent years. However, the very notion of ‘a’ 3D reconstruction is simplistic, genomes being dynamic and variable with differences according to organism, tissue, cell-type, cell-cycle, and cell. Hi-C experiments are frequently performed on large, synchronized cell-type specific populations, so that resultant reconstructions are interpreted as providing a consensus configuration. The emergence of single cell Hi-C (scHi-C, Ramani. et al., 2017; Stevens et al., 2017) has enabled dissection of inter-cellular structural variation, at the expense of yielding appreciably sparser data. Developing reconstruction methodology to accommodate such sparsity is one of our contributions; see Sections 11–14.

Another component of structural variation is allelic: in diploid organisms maternal and paternal homologs can adopt differing configurations. This poses difficulties for reconstruction algorithms since Hi-C readout is generally unphased, and resultant contacts are ambiguous as to whether they are intra- or inter-homolog. Until recently, these concerns have been ignored but novel approaches attempt to resolve identifiability issues by either prescribing assumptions and/or invoking additional data sources (Cauer et al., 2019; Belyaeva et al., 2021). We do not address these aspects here, but note that the concerns can be sidestepped if phasing of Hi-C output can be achieved, HiCHap (Luo et al., 2020) being an accurate tool for doing so. Clearly, the forefront structural feature of a single chromosome is its contiguity, followed by its folding complexity, necessary to achieve the compaction needed to fit within the nucleus. And it is these features that our original 3D reconstruction algorithm, Poisson Metric Scaling (PoisMS, Tuzhilina, Hastie and Segal, 2020) addressed. In prior work contiguity had been tackled by imposing constraints (Duan et al., 2010; Ay et al., 2014; Stevens et al., 2017), which are cell type specific and require prescription of constraint parameters. Given a paucity of relevant background biological measurement, these parameters can be difficult to specify. Further, their inclusion substantially increases computational burden. Other approaches (Zhang et al., 2013; Park and Lin, 2017; Rieber and Mahony, 2017) ignore contiguity in the reconstruction process, imposing it post hoc by “connecting the dots” of the 3D solution according to the ordering of corresponding genomic loci.

We review our PoisMS methodology, that extends principal curves (Hastie and Stuetzle, 1989) to the metric scaling problem, in Section 2. Previously, we had used B-spline bases as primitives for obtaining chromatin configuration but, as described in Section 3, this formulation is problematic with regard deploying cross-validation to determine smoothness degree (for reasons detailed in Sections 3 and 10), which is critical for appropriately capturing the abovementioned second key attribute, folding complexity. Accordingly, in Sections 3 through 9, we introduce a smoothing spline basis, and attendant algorithm SPoisMS, and describe how this improvement enables effective cross-validation, mitigates initialization concerns, can be efficiently implemented by exploiting connections with PoisMS which, in turn, it is shown to outperform in a series of experiments, and develop degrees-of-freedom estimates facilitating calibrated methods comparison.

We then turn attention to issues surrounding sparsity and overdispersion – characteristic of all contact matrices but especially extreme for data deriving from single cell Hi-C assays, the importance of which has already been noted. These features make Poisson assumptions inappropriate. To address these concerns we advance general distribution-based metric scaling methodology and algorithms (DBMS) (Section 12), then specialize to select special cases: hurdle Poisson, zero-inflated Poisson, negative binomial (Section 13), and finally make comparisons between these and the original Poisson formulation (Section 14) before providing concluding Discussion and directions for further work.

## 2. PoisMS method

We provide a brief overview of our recently proposed Poisson Metric Scaling (PoisMS, Tuzhilina, Hastie and Segal, 2020), a novel approach which directly models 3D chromatin configuration by a 1D smooth curve.

Suppose we are aiming to obtain the reconstruction of the chromatin with *n* genomic loci. Denote the matrix of the loci spacial coordinates by 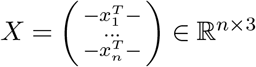. The core assumption of the model is

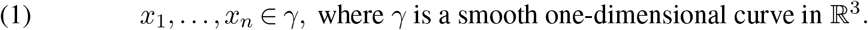

We start with transforming this assumption into a matrix form constraint as follows. First, if curve *γ* is parametrized by *t* and *t*_*i*_ index the genomic locus of *x*_*i*_ in the parameter space, then (1) can be reformulated as the set of equations *x*_*i*_ = *γ*(*t*_*i*_). Next, to incorporate the smoothing into the model, we claim that each component of *γ*(*t*) = (*γ*_1_(*t*), *γ*_2_(*t*)*, γ*_3_(*t*))^*T*^ is a cubic spline and decompose and, thus, can be decomposed in a linear combination of the basis functions. In other words, we state that

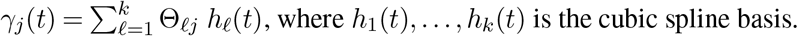

Here *k* is the size of the basis and is the hyperparameter that controls the “wiggliness” of the resulting reconstruction, which we will subsequently refer to as *degrees-of-freedom*. Finally, if *H* ∈ ℝ^*n*×*k*^ is the matrix with elements *H*_*iℓ*_ = *h*_*ℓ*_(*t*_*i*_) and Θ ∈ ℝ^*k*×3^ represents the spline coefficient matrix, then the smoothing constraint can be rewritten in matrix form as *X* = *H*Θ.

Next, we introduce the probabilistic model for the contact counts. To do so, we use Poisson distribution and link the Poisson parameters to the pairwise distances between loci. To be precise, if 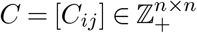 is the contact matrix then we assume that *C*_*ij*_ ~ Pois(*λ*_*ij*_) and

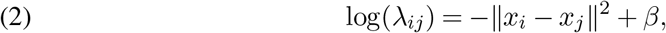

where *β* is some unknown intercept. The resulting negative log-likelihood objective is there-fore

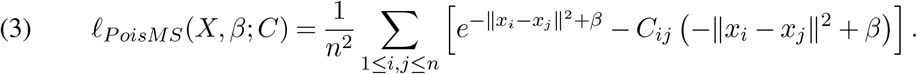

Combining (3) with the smoothing constraint leads us to the following MLE problem

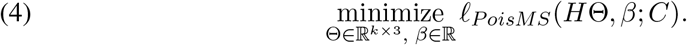

Note that the particular choice of the log-link (2) exploits the following heuristic. First, the loci that are close to each other in 3D space should have a higher expected contact frequency value. Second, although equivalent to the more traditional exponential link *λ*_*ij*_ = *β*‖*x*_*i*_ – *x*_*j*_‖^*α*^ proposed by, for example, Rosenthal et al. (2019), it results in the objective that is much easier to optimize. In Tuzhilina, Hastie and Segal (2020) we propose an efficient iterative algorithm that recovers the solution to (4), which, schematically, can be stated as follows.

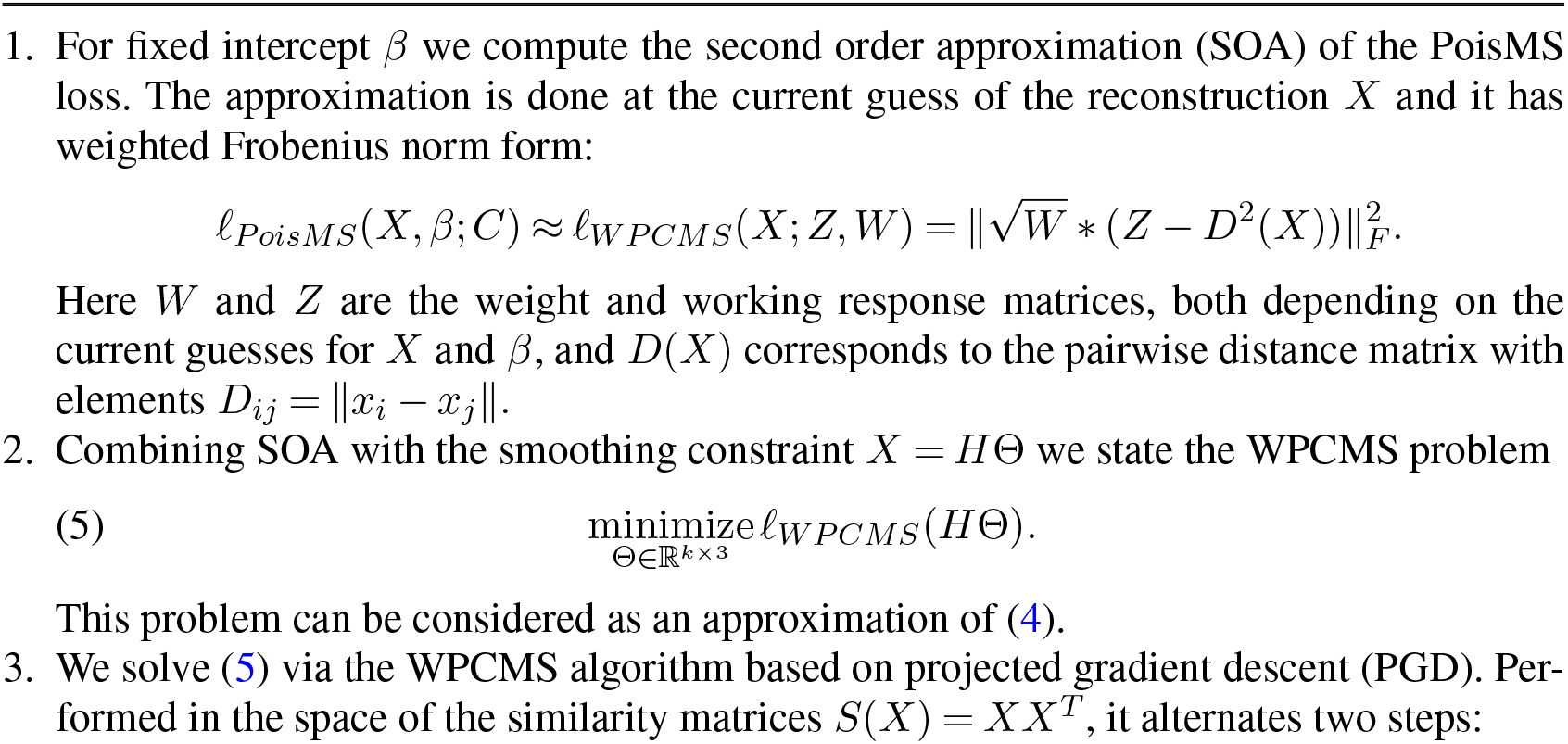
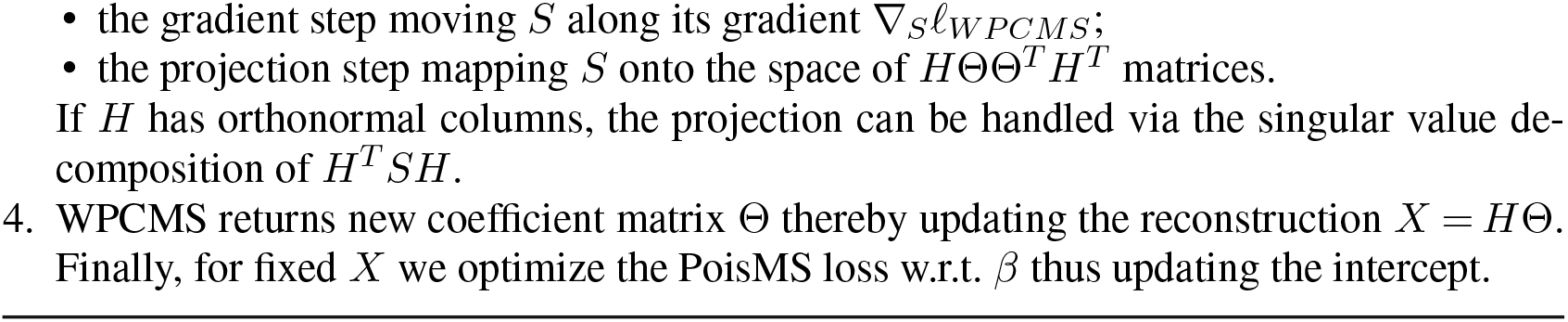
Poisson Metric Scaling (PoisMS)

## 3. Alternative approach: smoothing splines

There are a few caveats about the PoisMS model. First, since the objective in (4) is non-convex, the PoisMS algorithm converges to a local minimum. Thus the initialization for Θ and *β* can impact the resulting reconstruction. Although we did not observe significant variation in the PoisMS solution in our experiments, an initialization choice remains an open question. We provide more details in the Appendix of Tuzhilina, Hastie and Segal (2020). Second, it is not exactly clear how to use cross-validation in the context of the contact matrix. The original idea was to hold out some chromatin loci pretending that they were unobserved, train the PoisMS model on the observed loci and evaluate the fit on the held-out set. The training procedure is, therefore, equivalent to removing a subset of rows and columns in the contact matrix, eliminating the corresponding rows in the spline basis matrix, and reconstructing the part of the chromatin using only the observed blocks of *C* and *H*. Although such an approach may seem reasonable, one should be very careful selecting the unobserved set of loci. Due to the strong correlation in the contact matrix choosing the loci at random may result in inadequate cross-validation scores.

To demonstrate the correlation we take the IMR90 cell data for chromosome 20 and 100kb resolution, which contains *n* = 599 loci, and compute the reconstruction *X* via PoisMS. In doing so, we use 25 degrees-of-freedom, the value which was identified as the optimal in Tuzhilina, Hastie and Segal (2020). Next, we evaluate the matrix of the Poisson parameters 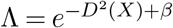 and compute the error matrix *E* = *C* – Λ. In Figure 1 we present the heatmap for the pairwise correlations between all the rows in *E*. According to the plot, in addition to the thick high correlation region along the diagonal, we observe multiple continuous off-diagonal areas, where the correlation is very closed to one. Both these patterns imply a strong correlation between the neighboring rows in the error matrix.

**FIG 1.**
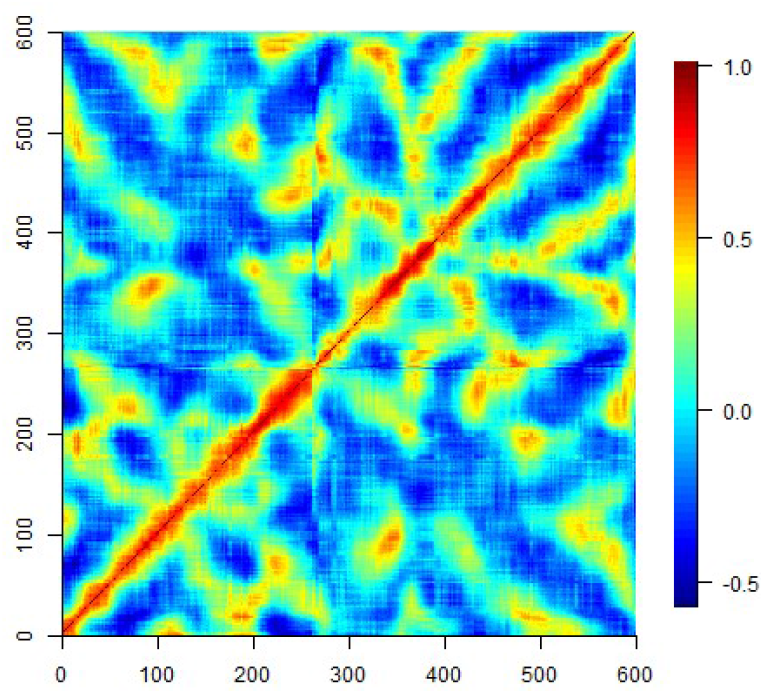
The heatmap of the correlation matrix measuring the association between the rows of the error matrix *E* = Λ – *C*. The reconstruction *X* is computed for *k* = 25 degrees-of-freedom. Since the diagonal elements of C are extremely large the error matrix has huge values on the diagonal as well. To mitigate the influence of the diagonal of E on the resulting correlation we measure the Spearman correlation instead. The thick red region along the diagonal and multiple red spots spread around the heatmap justify the strong correlation between the neighboring rows in the error matrix.

To counteract the interaction between the neighboring elements in the contact matrix, instead of selecting unobserved loci at random, we suggest using block cross-validation, thereby removing rows in *H* as well as rows and columns in *C* in continuous blocks. This is where PoisMS faces some difficulties. Specifically, since each element of the B-spline basis has finite support, the spline basis matrix tends to be sparse. Thus, eliminating a block of rows in *H* can potentially exclude one or multiple elements from the basis resulting in a very high reconstruction variance.

In this section we propose a novel approach based on the smoothing spline technique that resolves the initialization and cross-validation issued discussed above. First, we rewrite (3) in terms of the *γ* curve as

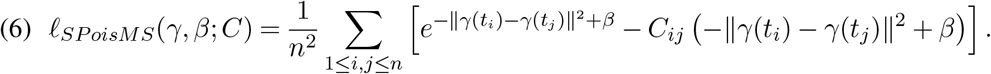

Instead of incorporating the smoothness through the B-spline basis we form the penalized loss

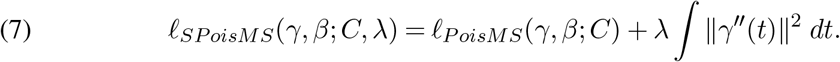

The second term in the objective measures the curve smoothness via its second-order derivative and is usually called *roughness penalty*. Therefore, we state an alternative *Smoothing*

*Poisson Metric Scaling (SPoisMS)* optimization problem

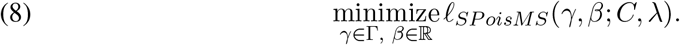

Here Γ is the set of all smooth (second-order derivative exists) one-dimensional curves in ℝ^3^. Note that in SPoisMS the “wiggliness” of the resulting reconstruction is controlled in a different way than in PoisMS, where it is done through the basis size *k*. In SpoisMS increasing *λ* will put more penalty on the second derivative of *γ* thereby encouraging more smoothness in the solution.

One can follow the standard smoothing spline theory to restate problem (8) in matrix form (see, for example, Green and Silverman (1994)). First, we claim that each component of the optimal solution *γ* is a natural cubic spline with knots *t*_1_,…, *t*_*n*_. Denote by *n*_1_(*t*),…, *n*_*n*_(*t*) the natural spline basis.

If *N* ∈ ℝ^*n*×*n*^ is the matrix representing the basis evaluations at knots, i.e. *N*_*iℓ*_ = *n*_*ℓ*_(*t*_*i*_), and Ω ∈ ℝ^*n*×*n*^ is the penalty matrix with elements 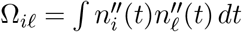, then for each *γ* there exists Θ ∈ ℝ^*n*×3^ such that the objective (7) is equivalent to

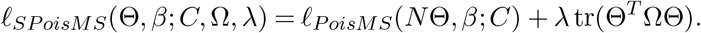

Now, let *K* = *N*^−*T*^ Ω*N*. Since *N* is full-rank and non-singular one can use the change of variables *X* = *N* Θ thereby restating the optimization problem (8) as

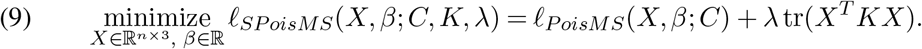

## 4. Link between PoisMS and the smoothing spline approach

Matrix *K* has several important properties. We will use these properties to demonstrate that the SPoisMS loss calculated for the original contact matrix *C* is equivalent to the PoisMS loss computed for 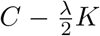. As a consequence, the SPoisMS problem can be easily solved by applying the original PoisMS method to adjusted version of the contact matrix.

Denote *K* = *UDU*^*T*^ the eigendecomposition of the penalty matrix. One can prove that *K* has two zero eigenvalues and the corresponding eigenvectors span the subspace of linear functions (see, for example, Green and Silverman (1994)). This fact can be used to validate the equations *K*1 = 0 and **1**^*T*^ *K* = 0, which further imply Σ_1≤*i,j*,≤*n*_ *K*_*ij*_ = 0 (see Appendix A for details). Here **1** = (1,…, 1)^*T*^ ∈ ℝ^*n*^ refers to the *n*-dimensional vector of ones. Using these properties we link the SpoisMS and PoisMS losses as stated below.

### 1. Lemma

*If K is the smoothing spline penalty matrix and* 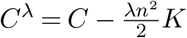 *then*

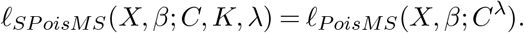

Proof. Recall that *S*(*X*) = *X X*^*T*^ represents the matrix of the pairwise inner products between genomic loci and *D*(*X*) corresponds to the pairwise distance matrix with elements *D*_*ij*_ = ‖*x*_*i*_ – *x*_*j*_‖. Denote by

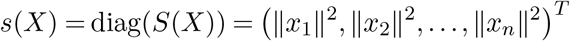

the diagonal of the inner product matrix. There is a link between *S*(*X*) and *D*(*X*), i.e.

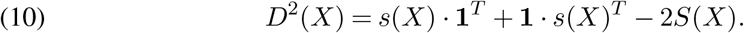

Combining this equation with the properties of *K* and the trace properties it is easy to show that

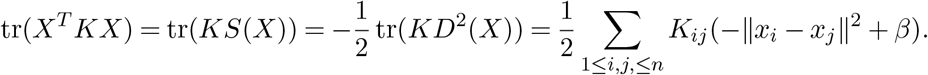

The following steps conclude the proof:

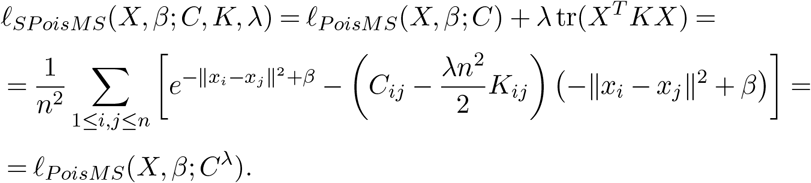

It should be noted that this lemma significantly simplifies the practical application of SPoisMS. Instead of developing new iterative algorithms that would solve (9), we could just use the original PoisMS technique on the adjusted contact matrix *C*^*λ*^.

## 5. Truncating the basis

Lemma 1 demonstrates that (9) is equivalent to solving the unconstrained PoisMS problem

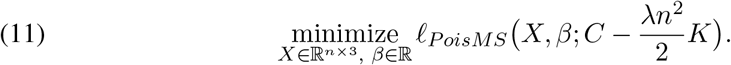

Recall that if there was the smoothing constraint *X* = *H*Θ, the projection step in the PoisMS algorithm would involve the singular value decomposition of the *k × k* matrix *H*^*T*^ *SH*, which is a cheap operation for small basis (see Section 2 for the details). However, in the case of the unconstrained problem (11) the projection becomes problematic as now it requires us to compute the SVD of the large *n × n* matrix *S* at each iteration.

There are several ways to get around this computational issue. Note that the PoisMS projection step aims to find Θ that minimizes the Frobenius distance between the similarity matrix *S* and *H*ΘΘ^*T*^ *H*^*T*^, which, for orthogonal *H*, is equivalent to solving

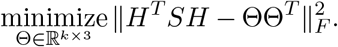

In other words, this step seeks for rank three approximation of *H*^*T*^ *SH*, so it is sufficient to compute only the first three singular vectors of this matrix. As the first computations trick, one can consider applying block power method (see, for example, Bentbib and Kanber (2017)), which can be more efficient for finding few singular vectors than calculating full SVD.

The second trick proposes to introduce a constraint to (11). It turns out that, for a proper constraint, the resulting problem would be an accurate approximation of (9) whereas requiring less costly SVD steps. Recall that we decomposed the penalty matrix as *K* = *U*^*T*^ *DU*. Since the columns of *U* form the natural spline basis (aka the *Demmler-Reinsch basis*), one can represent the resulting reconstruction as *X* = *U* Θ leading to an alternative form tr(Θ^*T*^ *D*Θ) for the smoothing penalty. Next, we refer to the following nice property of the Demmler-Reinsch basis: the wigglier is the basis element, the larger is the eigenvalue corresponding to it. In terms of the penalty, the smoothness in the reconstruction is encouraged by penalizing the most the coefficients in Θ corresponding to the largest diagonal elements in *D* and, as a result, related to the wiggliest part of the basis *U*. Since some of the *U* columns have a little effect on the resulting reconstruction we can remove them from the basis. In other words, we can set *H* ∈ ℝ^*n*×*k*^ to the eigenvectors from *U* corresponding to the largest *k* eigenvalues and solve the alternative problem

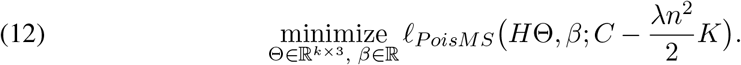

This can be easily done via the PoisMS approach again. For *k* large enough, the resulting solution will be an accurate approximation of the one from (11). However, now each projection step in PoisMS involves the SVD of smaller *k* × *k* matrix.

## 6. SPoisMS and PoisMS comparison

There are several advantages of using smoothing splines for the chromatin reconstruction.First, the PoisMS approach, where the size of the B-spline basis controls the reconstruction smoothness, requires the user to recompute matrix *H* for each value of degrees-of-freedom. In contrast, *H* is fixed in smoothing splines, and the penalty factor controls the model flexibility, thereby implying Θ ∈ ℝ^*k*×3^ of the same shape for any value of *λ*. This observation can be especially beneficial while choosing the PoisMS algorithm initialization. For instance, one can produce the *path of solutions* gradually decreasing the penalty factor and re-using Θ obtained for larger *λ* as a warm start for smaller values. Second, the support for each Demmler-Reinsch basis function is ℝ. Thus, returning to the cross-validation issues discussed in Section 3, block cross-validation would demonstrate more robust behavior comparing to the B-spline case.

The main disadvantage of the smoothing spline technique is that you need to keep the basis size *k* relatively large to get an accurate solution. This makes each step of the SPoisMS computationally more expensive than the PoisMS one. Further, in the case of PoisMS the flexibility of the resulting reconstruction is controlled by *k*, which corresponds to the B-spline basis size and which can be naturally interpreted as the degrees-of-freedom of the model. On the contrary, although it is obvious that increasing *λ* should decrease the degress-of-freedom for SPoisMS, it is hard to formally measure the *df* in this case. We tackle this problem in the following section.

## 7. Degrees-of-freedom

The notion of effective degrees-of-freedom is very well studied in the context of linear models and can be defined as the trace of the “hat” matrix (see, for example, Hastie, Tibshirani and Friedman (2009)). In this section we suggest a method for estimating *df* for the SPoisMS model. The general idea is to approximate the loss *ℓ*_*S PoisMS*_ by a quadratic function (equivalently, approximate the original problem by least squares) and use the definition developed for the linear regression.

### 2. Lemma

*Given some square matrix M* ∈ ℝ^*n*×*n*^ *denote by* Ф(*M*) *the operator that subtracts the row sums of M form its diagonal, i.e.* Ф(*M*) = *M* – diag(*M* **1**). *Suppose X*_0_ *is the resulting SPoisMS reconstruction. One can show that*

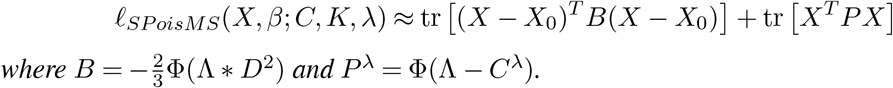

See Appendix B for the proof. Next recall that for the reduced Demmler-Reinsch basis we should add the constraint *X* = *H*Θ. Thus, the resulting approximation has a form of a weighted regression with a quadratic penalty, where the response corresponds to *X*_0_ and the feature matrix is *H*. Borrowing the definition we derive the formula for degrees-of-freedom

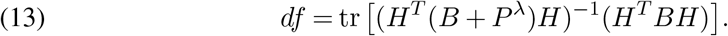

Before using (13), we slightly adjust it, so it follows two natural assumptions. First, if no penalty is imposed then degrees-of-freedom should be equal to the size of the basis used to produce the fit. In other words, for *λ* = 0 we expect *df* = rk(*H*), which can be achieved if *P*^0^ = Φ(Λ – *C*) = 0. Second, in the ideal setting, the degrees-of-freedom should not depend on the resulting reconstruction *X*. Both statements can be satisfied through replacing Λ with *C* or, equivalently, substituting expected counts with the observed ones. Note that, by analogy, one can replace *D*^2^ = *β* – log(Λ) by *β* – log(*C*), thereby implying the updated formulas

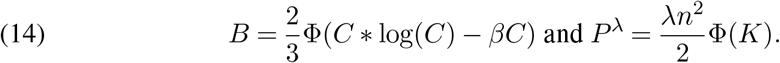

Here we assume that 0 . log(0) = 0 for zero contact counts.

## 8. IMR90 cell experiments

In this section we test the smoothing spline approach on the IMR90 cell chromosome 20 data.

First, we compute reconstructions via the SPoisMS method varying the penalty factor in the grid *λ* = 10^−4^, 10^−5^,…, 10^−9^. To speed up the computations we truncate the Demmler-Reinsch basis including only the eigen-vectors that correspond to the *k* = 300 smallest eigen-values of *K* (see Section 5 for more details). We produce the path of the solutions: we start from the largest *λ* = 10^−4^, and reuse Θ and *β* calculated at the previous step as a warm start for the subsequent value of *λ*. The resulting fits are presented in Figure 3. In the plot we observe that decreasing the penalty factor produces wigglier result as it imposes less smoothing penalty on the reconstruction. Next, for each *λ* value we report the corresponding degrees-of-freedom computed via (13) and (14), which we present in table below. According to the table, the considered range of *λ* covers degrees-of-freedom from *df* ≈ 6 (very smooth reconstruction) to *df* ≈ 117 (very wiggly). As a final step, to compare the performance of SPoisMS and the original method we round the resulting values of *df* and compute the PoisMS reconstructions using the B-spline basis of size *df* and random initialization. We demonstrate the resulting fits in Figure 3.

| $\lambda$ | $10^{-4}$ | $10^{-5}$ | $10^{-6}$ | $10^{-7}$ | $10^{-8}$ | $10^{-9}$ |
| --- | --- | --- | --- | --- | --- | --- |
| **df** | 6.15 | 11.32 | 20.73 | 37.56 | 67.07 | 117.43 |

**FIG 2.**
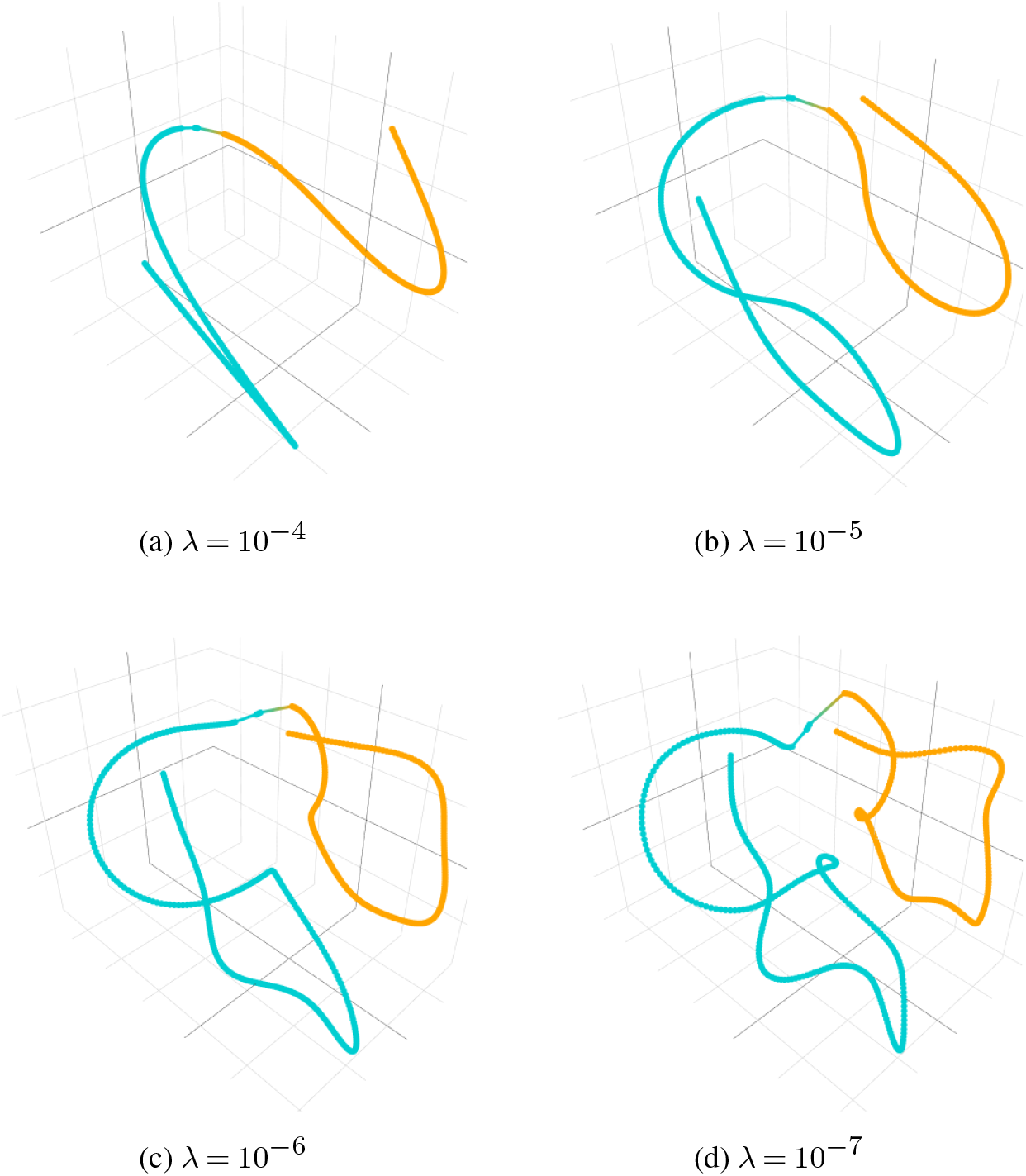
The projections of the resulting reconstructions obtained via the smoothing spline approach for different λ values. The solution path is produced in a sequential way: the reconstruction for larger λ is utilized as a warm start for the smaller λ. Colors (orange, teal) distinguish chromosome arms. Before potting the solutions were aligned via the Procrustes transformation.

**FIG 3.**
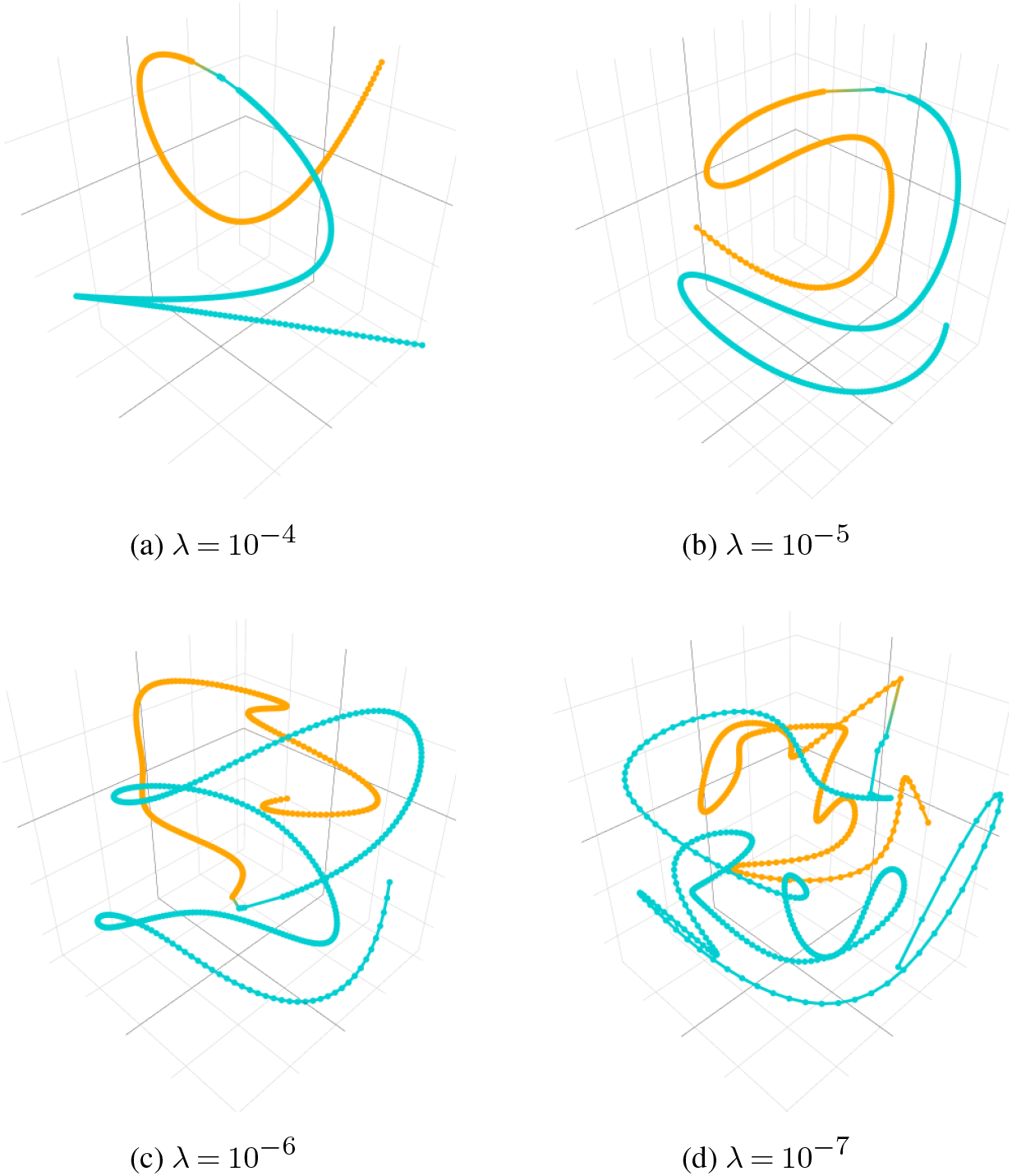
The projections of the resulting reconstructions obtained via the original PoisMS approach for different df values. The grid of df values is chosen to match the degrees-of-freedom in Figure 2. Each solution is produced by a random initialization of β and Θ. Colors (orange, teal) distinguish chromosome arms. Before potting the solutions were aligned via the Procrustes transformation.

Several conclusions could be drawn from Figures 2 and 3. First, comparing the PoisMS and SPoisMS solutions we observe that the degrees-of-freedom formula derived in Section 7 produces quite an accurate approximation of *df* for large *λ*, although it may underestimate *df* for small *λ*. Second, we conclude that reusing SPoisMS solutions as a warm start produces a well-aligned sequence of fits which allows us to track the evolution of the solution with the growth of *λ*. In contrast, we observe less agreement between the PoisMS solutions, which the random initialization of the method can explain. Finally, we notice that using warm-starts for subsequent *λ* values substantially decreases the number of iterations required for the method to converge. This illustrates the utility of the smoothing spline from the computational point of view.

## 9. Model validation

To further demonstrate the advantages of the SPoisMS approach we discuss how one can evaluate the reconstruction model using the contact matrix. In this section we propose an evaluation method based on a train-test split of genomic loci. Suppose we have only partial access to the contact matrix, i.e. we assume that some of the loci are unobserved. We denote by 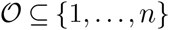 the set of observed loci and its complement by 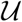. To assess our model we follow two steps:

1. We use the loci from 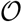 to compute the model coefficients Θ and intercept *β*.
2. Then we use 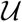 to evaluate the model’s performance.

In what follows, we explain how to tackle these two tasks for both the PoisMS and SPoisMS techniques.

### 9.1. PoisMS performance evaluation

To simplify the notations, without loss of generality, we can assume that the rows and columns in *C* are permuted in such a way that the first 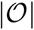 of them represent the contact counts for the observed loci. In other words, the contact matrix has block-structure 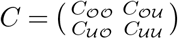, which, in turn, implies the structure in the basis matrix 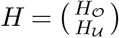.

We start with the PoisMS approach. To fit the model on observed data we evaluate the log-likelihood only at contact counts from 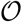 thereby leading to the train PoisMS loss

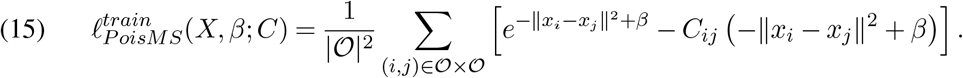

Here 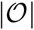 corresponds to the number of observed loci. It is not hard to show that

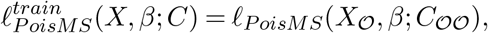

therefore, minimizing (15) subject to the smoothing constraint *X* = *H*Θ is equivalent to solving the problem

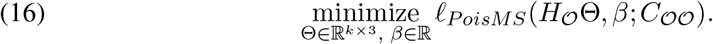

In other words, the solution can be found by running PoisMS for the contact matrix 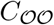 and basis 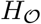.

Now we proceed to the model evaluation step. Note that the original PoisMS algorithm requires the basis matrix to have orthonormal columns, which is true for the full *H*, but may not be the case for the sub-matrix 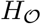. Thus before fitting the PoisMS model we need to compute the *QR*-decomposition 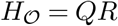 and replace the basis matrix by *Q*. Solving the PoisMS problem results in the coefficient matrix Θ as well as the intercept *β*. In order to evaluate how well these parameters fit the remaining contact counts we need to compute the complementary test loss

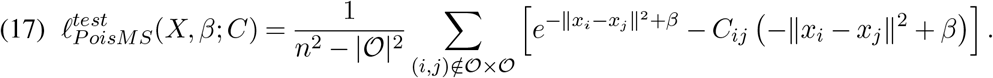

Since this loss involves the contacts between both observed and unobserved genomic loci it is necessary to recover the full reconstruction, which can be easily obtained via the formula

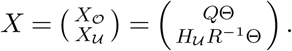

This reconstruction will be subsequently plugged in (17) thus producing the model test score.

### 9.2. SPoisMS performance evaluation

Now we extend the approach proposed in the previous section to the smoothing spline technique. By analogy with (15), one can derive the SPoisMS train loss as follows

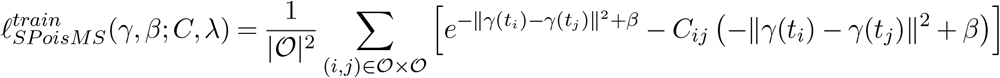

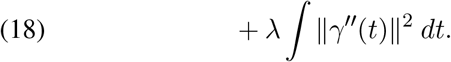

For the penalized loss the following lemma can be proved.

#### 3. Lemma

*The optimization problem*

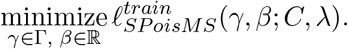

*is equivalent to solving the reduced PoisMS problem*

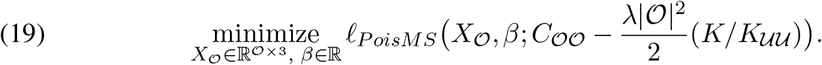

*Here 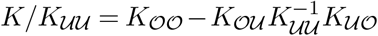 is the Schur complement of the penalty matrix, which has block structure similar to C, i.e.* 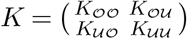.

The proof is based on the properties of the smoothing spline basis (see Appendix C for the details). The lemma demonstrates that the solution to the SPoisMS problem can be computed by means of the PoisMS algorithm, even for a subset of knots. Moreover, it implies that if we treat some loci as unobserved one can re-use full *K* to compute the penalty matrix for the reduced problem. In the lemma proof we also presented the explicit formula to impute the unobserved part of the reconstruction. The complete reconstruction

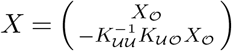

will be subsequently plugged in (17) thereby producing the test score.

Note that the idea form Section 5 can be extended to the partially observed data as well. Specifically, to reduce the computations of the train fit one can obtain an accurate approximation of (24) by taking the SVD of 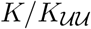 and using a subset of singular vectors as a basis for 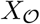.

## 10. Block cross-validation

In Section 3 we raise the issue of high-correlation between the contact counts, which makes the standard cross-validation approach problematic in the context of the chromatin reconstruction. To moderate the influence of correlated data on the model evaluation process, we suggest to use block cross-validation (BCV). In this section we investigate the BCV performance of the PoisMS and SPoisMS approaches using the IMR90 cell data.

First, we split *n* = 599 genomic loci into 20 folds, each containing consecutive loci. We assign one fold to 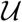 thereby treating 5% of loci as unobserved; the remaining 19 folds, or 95% of loci, are assigned to 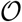. Next, following the procedure from Section 9.1, we use 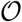 to compute the solution (Θ, *β*) either via PoisMS of via SPoisMS. In our experiments, we test SPoiMS with different penalty factors *λ* = 10^−4^, 10^−5^,…, 10^−9^. To make the PoisMS results compatible to the SPoisMS ones, we use the degrees-of-freedom from Table **??** when computing the PoisMS solution (see Section 8 for the details). As a result, for each of the method and each of the hyperparameter we obtain the coefficient matrix Θ and the intercept *β*. Finally, to evaluate how well these parameters fit the unobserved set of loci 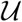 we impute the reconstruction and use the completed *X* to evaluate the test score (17). We repeat the entire experiment for all folds except the boundary ones and present the mean as well as the 1SE intervals of the test score in Figure 4.

**FIG 4.**
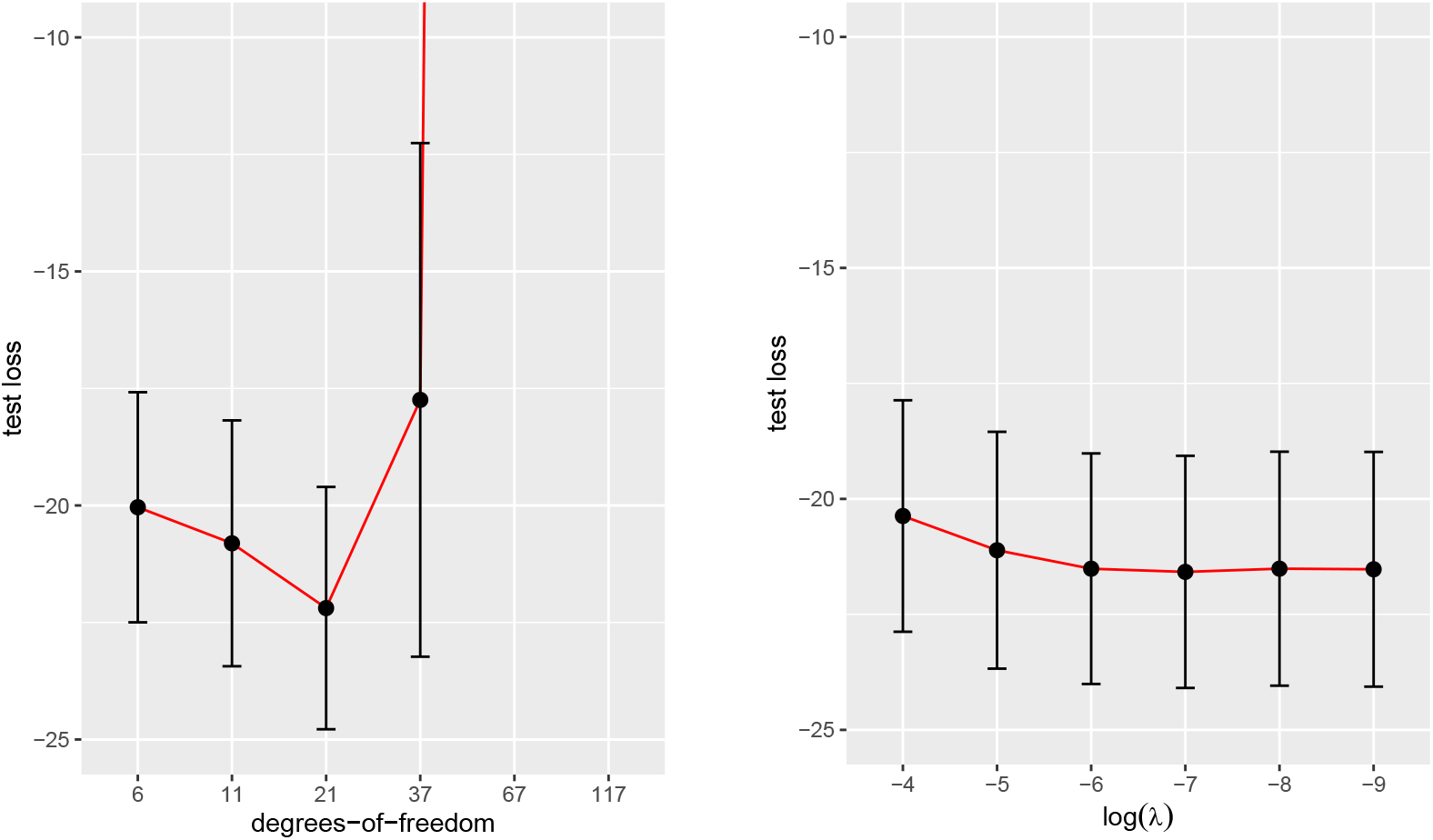
Comparison of the block cross-validation performance for the PoisMS and SPoisMS methods. Left panel: the test score is computed via PoisMS with different degrees-of-freedom. For df = 37 the confidence interval is much wider than for small degrees-of-freedom, for df = 67 and 117 the mean test score is so large that the confidence interval did not fit in the plot limits. Right panel: the test score is computed via SPoisMS for a grid of penalty factors (aligned with the df values from the left panel). The train fit is obtained using 95% of the loci, the test score is computed on the remaining 5% of the loci.

From the left panel of the plot we infer that, as expected, the confidence intervals computed via PoisMS “explode” for high values of degrees-of-freedom. On the contrary, the SPoisMS test score exhibits robust behavior regardless of the penalty factor *λ*. To summarise, introducing smoothness to PoisMS through the roughness penalty instead of the B-spline basis finally enables reconstruction evaluation by means of block cross-validation.

## 11. Sparse contact matrices and over-dispersion

The contact matrix has two important features. First, usually is it very sparse, i.e. contains big portion of zero counts. Second, it is diagonally dominant (Yang et al., 2017). Both of these factors may cause zero-inflation and over-dispersion of the data thereby making the Poisson distribution unsuitable for modeling the contact counts. To check if this is the case for the IMR90 data we take the reconstruction *X* produced by the PoisMS approach with *df* = 25 and compute the expected mean count matrix Λ (see Section 3 for more details). In Figure 5 we present the scatter plot for the observed counts *C*_*ij*_ vs. the expected ones Λ_*ij*_. The dense conglomeration of the points along the x-axis of the figure as well as the significant deviation of the observed values from the predicted ones for high contact counts reveals the potential presence of zero-inflation and over-dispersion in the data.

**FIG 5.**
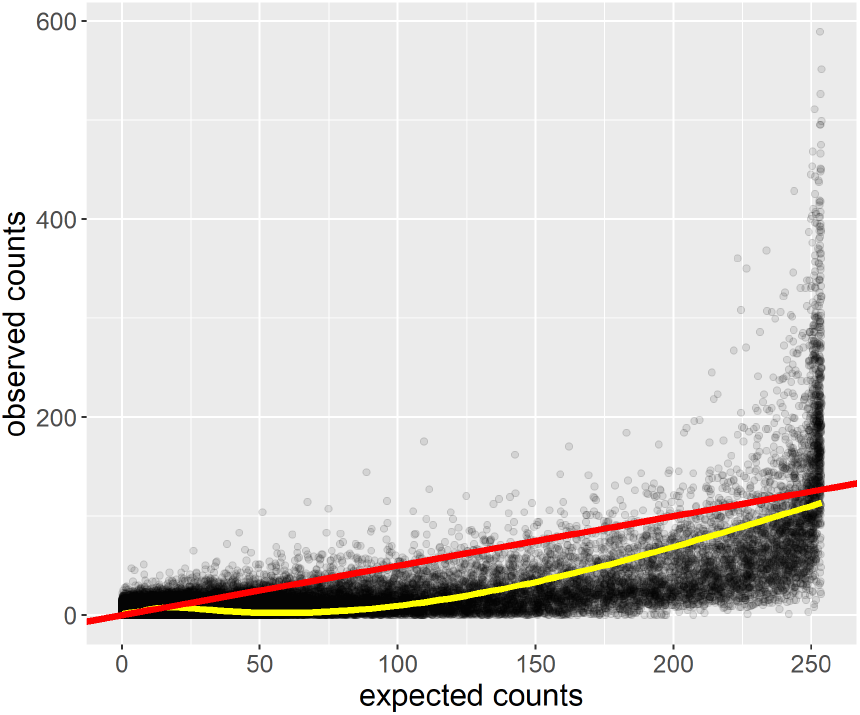
Observed vs. expected counts. In this plot we present the dependence between the contact counts *C*_*ij*_ and the Poisson parameters Λ_*ij*_ obtained by PoisMS with *k* = 25 degrees-of-freedom. The reconstruction is computed for IMR90 data. Red line: represents the ideal fit where observed and expected counts are equal. Yellow line: represents the smooth approximation of the scatter plot; the zero-inflation of the contact counts is demonstrated by its clinging to the expected counts axis.

## 12. Distribution-based metric scaling

The main concern for using Poisson distribution when modeling the contact counts is that the resulting reconstruction is highly influenced by abundant zeroes and the diagonal of the contact matrix. To mitigate the influence of these data artifacts, one can consider alternative distributions such as zero-inflated Poisson or negative binomial. It turns out that one can easily extend the PoisMS algorithm outline from Section 2 to other distributions as well. Below we propose general *distribution-based metric scaling (DBMS)* and demonstrate how one can use this technique to build three alternative chromatin reconstruction models.

The DBMS model assumes that each contact count follows a discrete distribution with support ℤ_+_. This distribution depends on some parameters, which we subsequently link to the chromatin conformation *X*. We propose that the link function involve pairwise distances between loci as well as some nuisance parameters that we denote by Ω. In other words, the resulting negative log-likelihood loss has the form of *ℓ*(*X*, Ω; *C*). To solve the DBMS problem we minimize the loss w.r.t. unknown *X* and Ω and subject to the smoothing constraint *X* = *H*Θ. Similar to the PoisMS case, the optimization algorithm alternates two steps.

### Step 1: update the reconstruction

Since the negative log-likelihood depends on the distances only, we can re-write the loss in the form *ℓ*(*D*^2^(*X*), Ω; *C*). As a first s tep, f or the current reconstruction guess *X*, we compute the second order approximation of the loss at the point *D*^2^(*X*). To do so, we calculate the first and second order derivatives

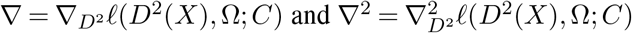

and evaluate the weights as well as the working response matrices

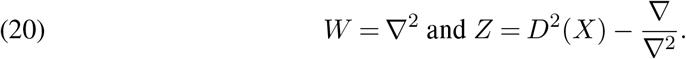

This leads us to the second order approximation

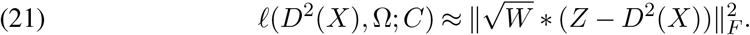

Now, instead of optimizing the original loss we minimize its second order approximation subject to the smoothing constraint *X* = *H*Θ. In other words, we solve the problem

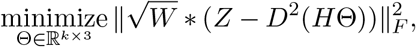

which can be done via WPCMS. As a result, we update Θ and the reconstruction *X* = *H*Θ.

### Step 2: update the nuisance parameters

We fix *X* and optimize the loss *ℓ*(*D*^2^(*X*), Ω; *C*) w.r.t. the Ω. Depending on the distribution, we will use one of the following approaches to update the nuisance parameters:

- compute the first order derivative of the loss w.r.t. Ω and use it to find an explicit formula for the optimal value;
- compute the first and second order derivatives of the loss w.r.t. Ω and use them to run Newton’s algorithm.

## 13. Alternative distributions

Below we suggest three alternative distributions for the contact counts: Hurdle Poisson, zero-inflated Poisson, and negative binomial. The optimization algorithm for each of them can be easily built using the DBMS outline from Section 12, which we discuss in detail in the Appendix.

### 13.1. Hurdle Poisson

Modeling the contact counts with Hurdle Poisson consist of two parts (see, for example, Gurmu (1998)). We first assume that attaining zero count follows the Bernoulli distribution; we further introduce the zero-truncated Poisson (ZTP) distribution for non-zero counts, i.e.

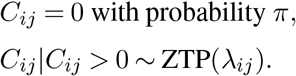

This leads us to the following distribution of *C*_*ij*_

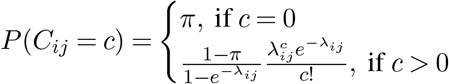

We again use the log-link (2) to introduce the dependence between the truncated Poisson parameters *λ*_*ij*_ and the chromatin 3D spacial structure. If 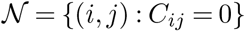 represents the subset of zero contact counts, then the goal of the *Hurdle Poisson metric scaling (HPoisMS)* is to minimize the negative log-likelihood

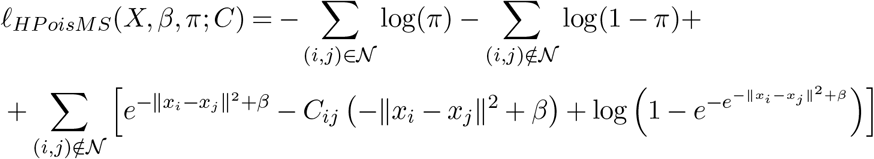

The optimal solution can be found via the HPoisMS algorithm that we discuss in Appendix D.1.

### 13.2. Zero Inflated Poison

The zero-inflated Poisson distribution addresses the abundance of zero contact in a different way (Lambert, 1992). It assumes that zero observations have two different origins: “structural” and “sampling”. The structural zeros occur with some fixed probability *π*, whereas the sampling zeros appear as a null Poisson outcome, i.e.

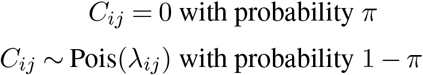

This leads us to the following distribution of the contact counts

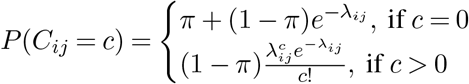

which, combined with the same log-link (2), implies the corresponding loss function

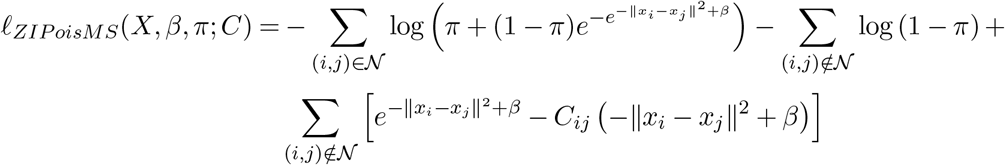

Zero-inflated Poisson metric scaling (ZIPoisMS), an optimization procedure that finds the optimal *X, β* nd *π*, is proposed in Appendix D.2.

### 13.3. Negative Binomial

Next, we consider the negative binomial model. We use the following distribution for the contact frequencies:

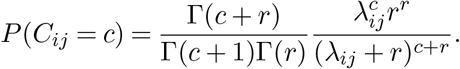

Comparing to the Poisson approach, which implies E(*C*_*ij*_) = Var(*C*_*ij*_), the negative binomial technique links the contact count mean and variance through the relation 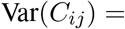 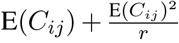. The nuisance parameter *r* adds more flexibility to the model thereby allowing overdispersion to be addressed.

Linking parameter *λ*_*ij*_ to the pair-wise distances ‖*x*_*i*_ – *x*_*j*_‖ through equation (2) one can state the goal of *negative binomial metric scaling (NBMS)*: find the optimal *X, β, r* that minimize the negative log-likelihood

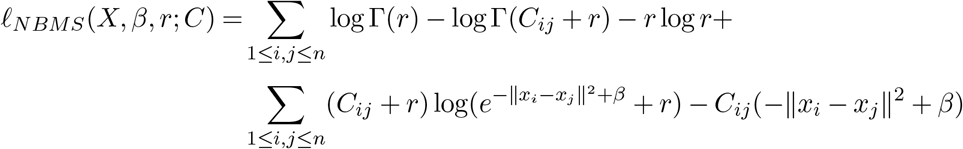

See Appendix D.3 for the details on the NBMS optimization algorithm.

## 14. Four models comparison

In this section we model contact counts by means of the four distributions discussed above (Poisson, Hurdle and zero-inflated Poisson as well as negative binomial). We start with a simple visual comparison of the models. We use IMR90 data and obtain four reconstructions approximation the chromatin curve by a B-spline with 25 degrees-of-freedom. Comparing the images in Figures 6, we can conclude that the Negative Binomial model tends to separate before and after centromere parts of the chromatin (blue and orange color in the plot). In contrast, the models that account for excessive zero counts, i.e. Hurdle and zero-inflated Poisson, imply more interaction between these two parts. This observation is also supported by Figure 7, where we present the heatmap for Λ(*X*) = *e*^−*D*(*X*)+*β*^. In the NB panel we clearly see two prominent diagonal blocks evidencing strong interaction within each of the two pieces of chromatin; the two off-diagonal blocks, however, have values closed to zero and imply scarce cross-contact. From the same plot one can infer that, unlike HPoisMS and ZIPoisMS, the NBPoisMS model was able to moderate the influence of the diagonal elements of the contact matrix on the resulting reconstruction.

**FIG 6.**
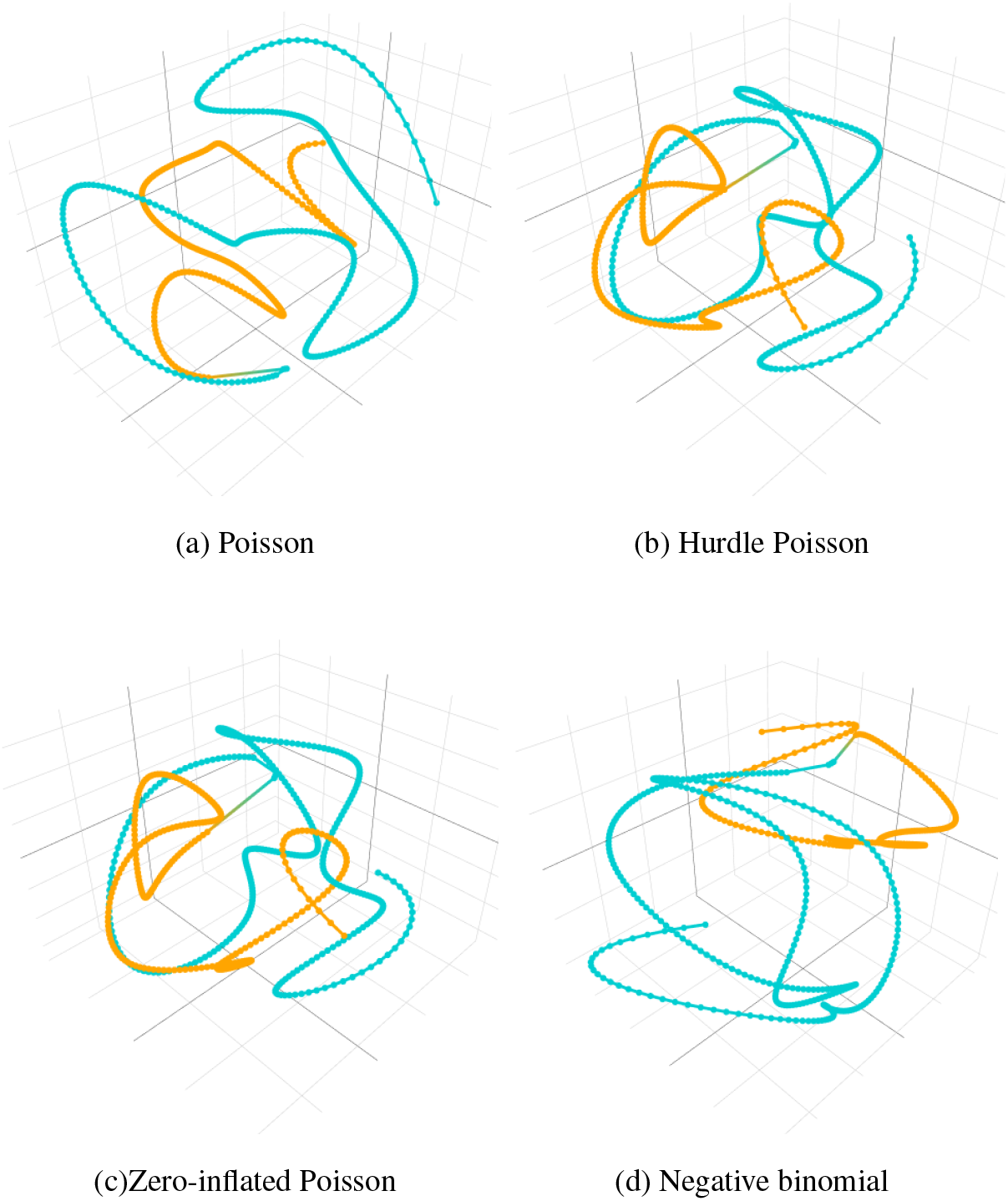
The projections of the resulting reconstructions obtained via four methods: PoisMS, HPoisMS, ZIPoisMS, NBMB. The degrees-of-freedom is set to 25 for the B-spline basis.

**FIG 7.**
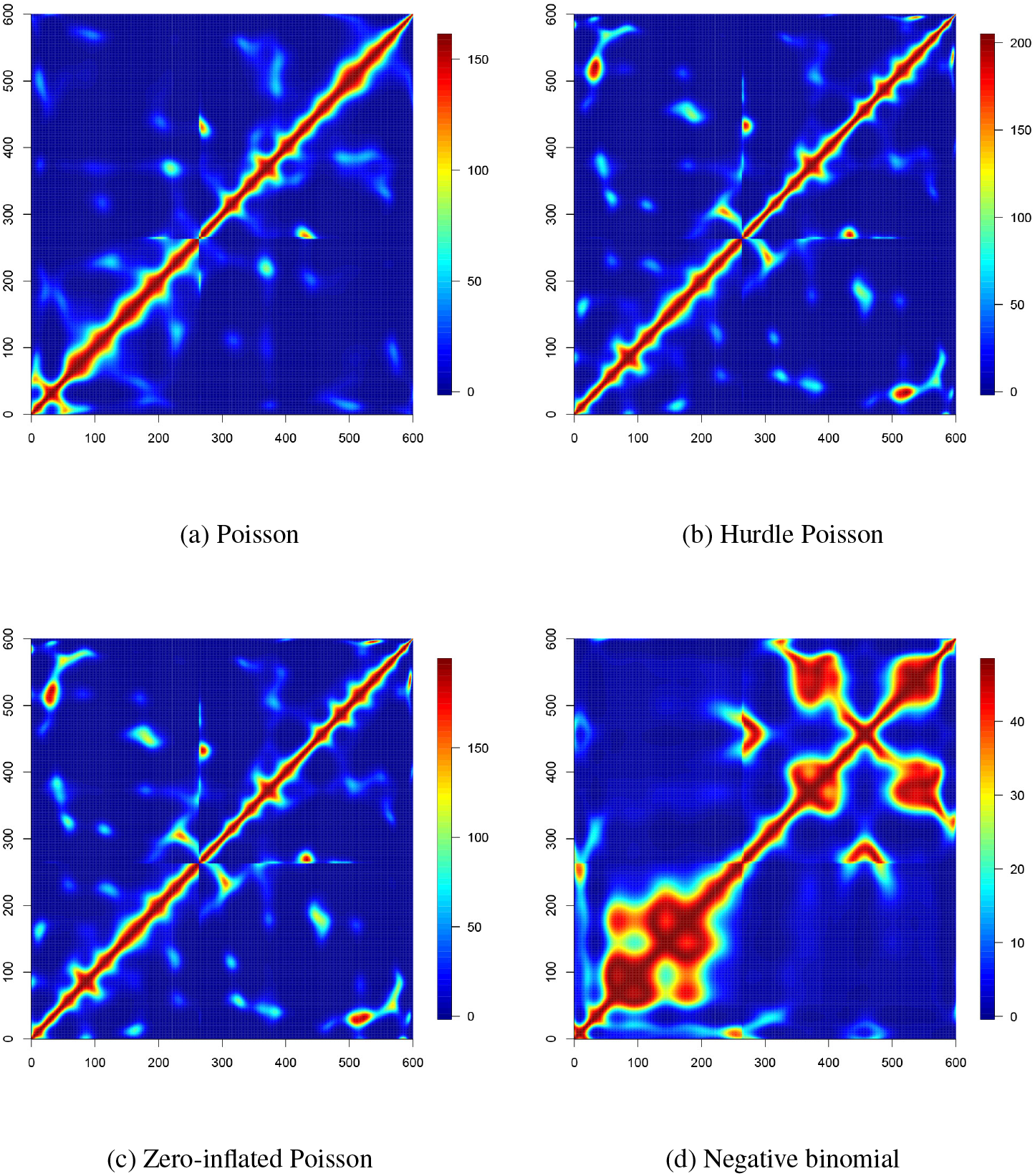
The heatmaps for Λ = *e*^−*D*(*X*)+*β*^ the resulting reconstructions obtained via four methods: PoisMS, HPoisMS, ZIPoisMS, NBMB. The degrees-of-freedom is set to 25 for the B-spline basis.

To compare the models performance, we use contact matrices for chromosome 1 from eight single mouse embryonic stem cells (Stevens et al., 2017, Gene Expression Ominbus (GEO) repository GSE80280) denoted *C*^(1)^,…, *C*^(8)^. Using data at 100kb resolution results in *n* = 1924 genomic loci. We run the following experiment. We select at random a subset four indices 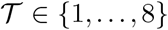 and calculate train and test contact matrices as 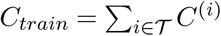 and 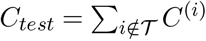 We use *C*_*train*_ to fit a model, thereby obtaining the reconstruction *X* and the optimal nuisance parameters. We subsequently use *C*_*test*_ to evaluate the model’s test score represented by the negative log-likelihood value. We repeat this experiment *N* = 30 times and calculate the average test score (across the random splits) as well as 1SE confidence interval.

In Figure 8 we plot the dependence of test performance on degrees-of-freedom value for each of the four models. Comparing the PoisMS and NBMS curves (blue and green), we conclude that using Negative Binomial distribution for contact counts in place of Poisson does not enhance the conformation reconstruction. On the other hand, we observe a substantial improvement in the test score for the other two models, where the best log-likelihood value is attained by ZIPoisMS with *df* = 10. The extreme sparsity of the contact matrix can explain such a difference in performance: the data has more than 98% of zero contact counts, thus, it strongly benefits from zero-inflated models.

**FIG 8.**
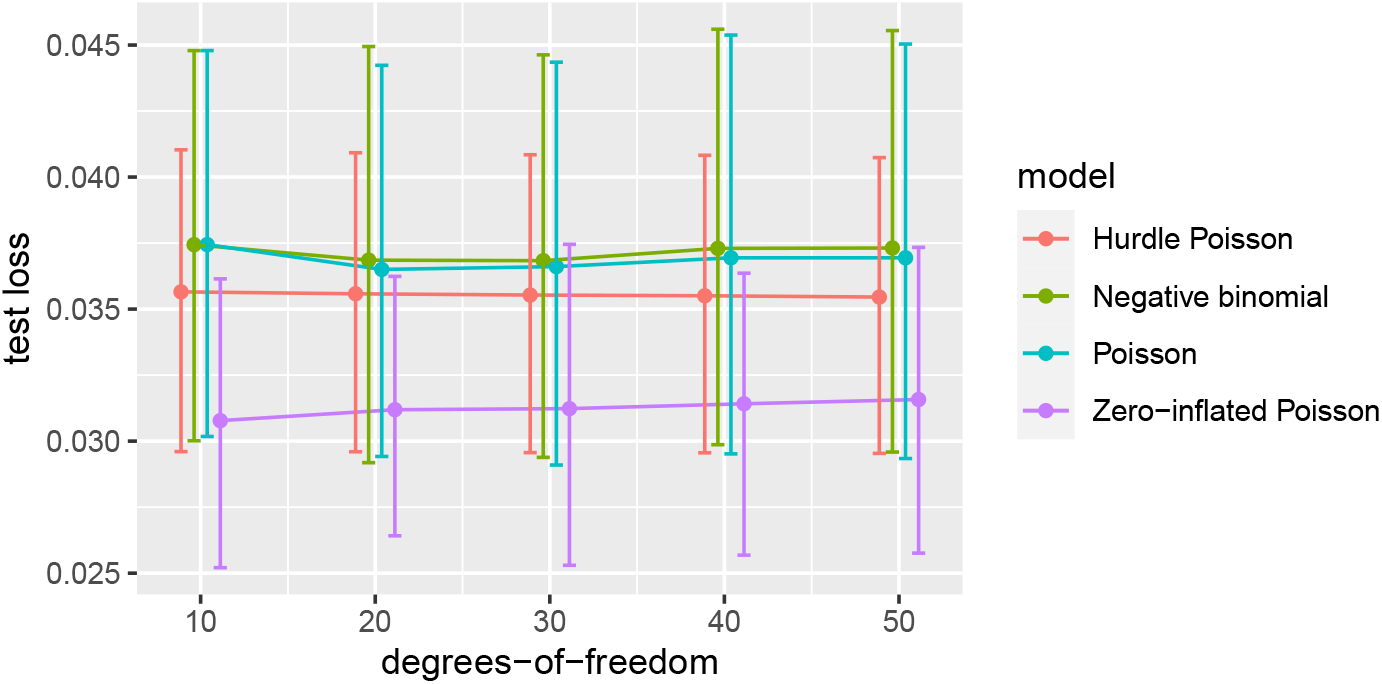
Comparing four models via the single cell data. Each model was fitted o n C train a nd subsequently evaluated on Ctest. The plot represents the average (across different train-test splits) test score vs degrees-of-freedom; the intervals correspond to the 1SE bands of the test scores. The plot evidences the superior performance of the ZIPoisMS models to the competitors.

## 15. Discussion

In this paper we propose several extensions to the PoisMS methodology described in Tuzhilina, Hastie and Segal (2020). First, we suggest an alternative way to encourage the reconstruction smoothness. Specifically, instead of using constrain (1), we combine the PoisMS objective with the roughness penalty. The resulting SPoisMS optimization problem (7) is a blend of PoisMS and smoothing splines, and, as we prove in Section 4, it has the following nice property: the reconstruction can be found by means of the original PoisMS algorithm. The motivation for such an alternative smoothing technique was the incompatibility of B-splines with block cross-validation and the inability to re-use the solution for smaller degrees-of-freedom as a warm start for the larger ones. We demonstrate the advantages of SPoisMS via IMR90 data set the in Section 10.

Next, we extend PoisMS in a different direction. We propose three alternative models for the contact counts: zero-inflated a nd H urdle P oisson as w ell a s N egative Binomial. These distributions were motivated by various artifacts present in the Hi-C data such as sparsity or diagonal dominance of the contact matrix. In section 12 we introduce a general framework that allows us to build optimization algorithm for a wide class of distributions. We subsequently use it for developing the ZIPoisMS, HPoisMS and NBMS techniques, which we finally compare via the single cell data.

There is still much scope for future experiments. First, we plan to test the novel techniques and validate our findings on data sets involving other chromosomes and resolutions. From a methodological point of view, we aim to enrich the class of models. In particular, we will explore Hurdle Negative Binomial distribution, which would simultaneously address sparsity and overdispersion. Moreover, we can refine the methods proposed in this paper by linking the nuisance parameters to the chromatin spatial structure. As an example, in the Hurdle model one can replace the common parameter *π* by *π*_*ij*_, different for each pair of loci, which we will link to the pair-wise distances via the logit link 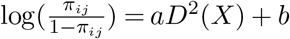.

Another interesting and more practical research direction would be to develop a more efficient implementation of the proposed algorithms. For instance, various acceleration techniques such as Nesterov and Anderson acceleration are known to perform well when combined with the projected gradient descent (see, for example, Tuzhilina and Hastie (2021)). Thus, these techniques can significantly speed up the convergence of the main building block WPCMS thereby enabling much faster computations for the DBMS solutions. Finally, the sparsity of the contact matrix can be an important feature used for improving the storage and computation time.

## Funding

E.T. was supported by Stanford Data Science scholarship. T.H. was partially supported by grants DMS-1407548 and IIS 1837931 from the National Science Foundation, and grant 5R01 EB 001988-21 from the National Institutes of Health. M.S. was partially supported by grant GM-109457 from the National Institutes of Health.

## SUPPLEMENTARY MATERIAL

### Code

Proposed methods are implemented in the R package DBMS; the software is available from Github (https://github.com/ElenaTuzhilina/DBMS).

## APPENDIX A PROPERTY OF THE DEMMLER-REINSCH PENALTY MATRIX

### 1. Lemma

*If* **1** = (1,…, 1)^*T*^ ∈ ℝ^*n*^ *is n-dimensional vector of ones and K is the smoothing spline penalty matrix, then*

- *K***1** = 0;
- **1**^*T*^ *K* = 0;
- Σ_1≤*i,j*,≤*n*_ *K*_*ij*_ = **1**^*T*^ *K***1** = 0.

Proof. Denote *K* = *UDU*^*T*^ the eigendecomposition of the penalty matrix. One can show that *K* has two zero eigenvalues and the corresponding eigenvectors span the sub-space of linear functions (see, for example, Green and Silverman (1994)). For simplicity, we assume that the eigenvalues are sorted in increasing order in *D*, so *d*_1_ = *d*_2_ = 0 and matrix *U* has the following structure: 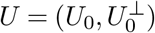, where *U*_0_ ∈ ℝ^*n*×2^ corresponds to the null-space of *K* and 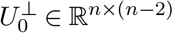 is the orthogonal space. Therefore, if *g*(*t*) is some linear function of *t* and *G* = (*g*(*t*_1_),…, *g*(*t*_*n*_))^*T*^, we get *G* ∈ *span*(*U*_0_) as well as 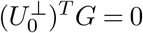. This leads us to the relation

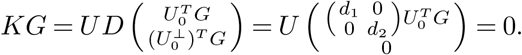

In particular, setting *g*(*t*) = 1 implies *KG* = *K***1** = 0. The second relation **1**^*T*^ *K* = 0 automatically follows form the fact that *K* is positive semi-definite (PSD), which immediately implies Σ_1≤*i,j*,≤*n*_ *K*_*ij*_ = **1**^*T*^ *K***1** = 0.

## APPENDIX B DEGREES-OF-FREEDOM

### 2. Lemma

*Given some square matrix M* ∈ ℝ^*n*×*n*^ *denote by* Φ(*M*) *the operator that subtracts the row sums of M form its diagonal, i.e.* Φ(*M*) = *M* – diag(*M* **1**). *Suppose X*_0_ *is the resulting SPoisMS reconstruction. One can show that*

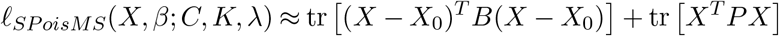

*where* 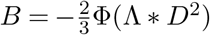 *and P*^*λ*^ = Φ(Λ – *C*^*λ*^).

Proof. Denote by *A*^*(ij)*^ the matrix such that all the elements are equal to zero except

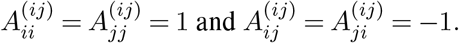

If *i* = *j* we can assume that *A*^*(ij)*^ = 0. One can show that for any matrix *M* ∈ ℝ^*n*×*n*^ the following relation holds

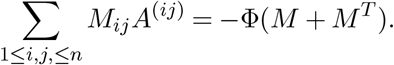

Further, combining

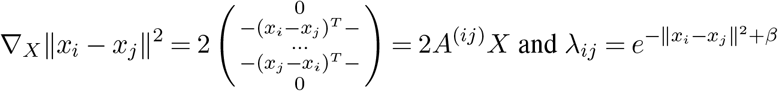

we derive that ∇_*X*_*λ*_*ij*_ = −2*λ*_*ij*_*A*^(*ij*)^*X*. Thus we compute the first order derivative of (3) as

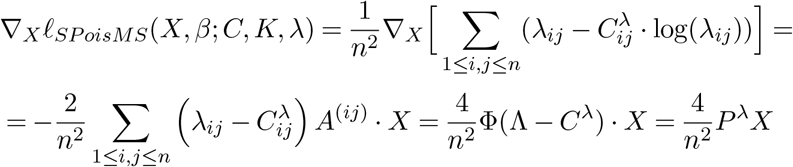

Here we exploit the fact that both Λ and *C* are symmetric matrices. Next we compute the second order derivatives. To make it simpler we compute them with respect to each column in *X* separately. Denote the columns of *X* by *X*_*i*_, i.e. *X* = (*X*_1_, *X*_2_, *X*_3_). If *f*: ℝ^*n*×3^ → ℝ is some arbitrary scalar function, then

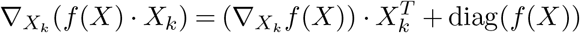

From the first derivative equations it follows that

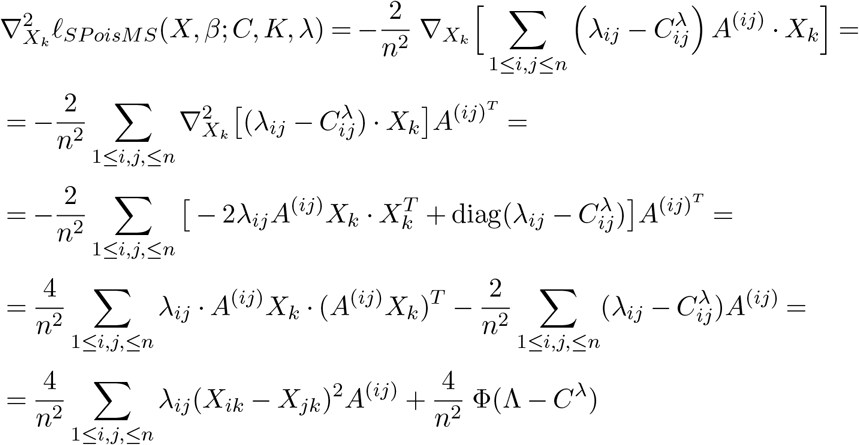

To make it simple we will approximate the actual Hessian by the average of the second order derivatives, which has a nice closed form

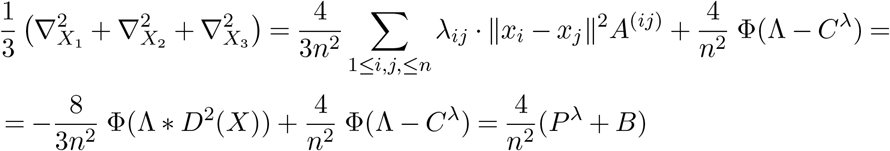

This leads us to the approximation (up to a constant) as follows

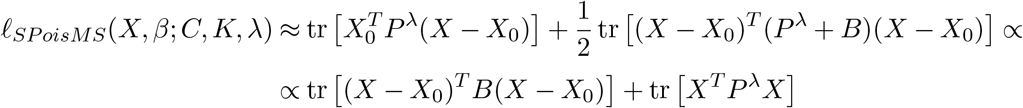

Note that the second order approximation in the original form should include terms 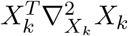 as well as the cross-terms 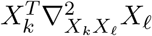, where 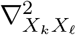 is the mixed second derivative computed w.r.t. both *X*_*k*_ and *X*_*ℓ*_. In our case we ignored the mixed terms and replaced all 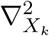 by their average. In other words, we approximate the original Hessian by a block-diagonal version with the same block 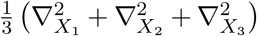 on the diagonal.

## APPENDIX C SPOISMS PERFORMANCE EVALUATION

### 3. Lemma

*The optimization problem*

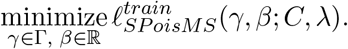

*is equivalent to solving the reduced PoisMS problem*

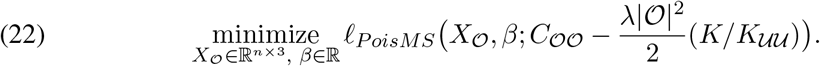

*Here* 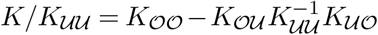 *is the Schur complement of the penalty matrix, which has block structure similar to C, i.e.* 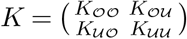.

Proof. First we note that Denote the solution by 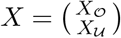. We begin with rewriting the loss function in matrix form as

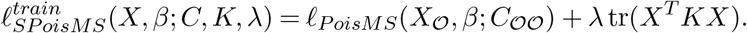

Next, we notice that the first part of the loss does not depend on 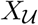. If 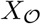 is fixed then 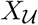 can be found as a minimum of

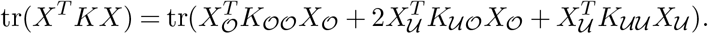

Taking the derivative w.r.t. 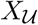 and setting it to zero leads us to the stationary point 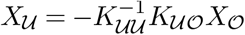. Plugging it back to the original loss function implies

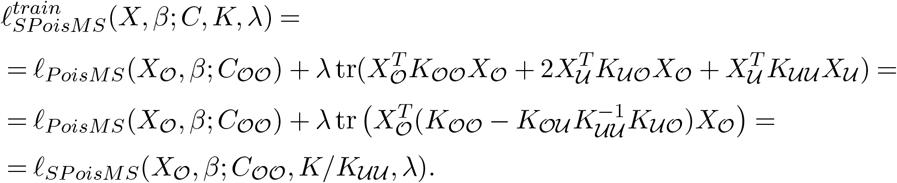

Therefore, the optimal 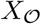 can be found by solving the SPoisMS problem for the contact matrix 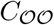 and the penalty matrix 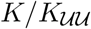, i.e.

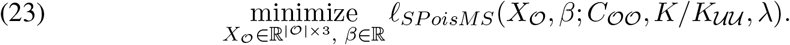

Note that in the main paper we demonstrated that the SPoisMS problem with the Demmler-Reinsch penalty matrix can be efficiently solved via the PoisMS algorithm. We extend this result to the optimization (23) as well. From the penalty matrix property *K***1** = 0 one can derive the system on equations involving the blocks of *K*, i.e. 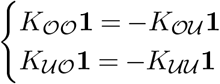, leading us to analogous property of the Schur complement

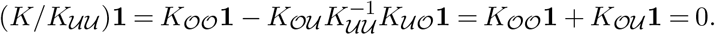

Following the proof in the main paper, one can show that the smoothing penalty in the loss function 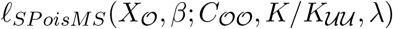 can be absorbed into the PoisMS part thereby proving the equivalence between (23) and the PoisMS problem

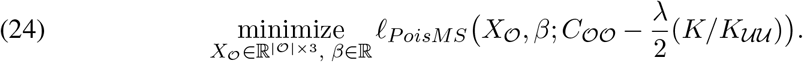

## APPENDIX D DISTRIBUTION BASED METRIC SCALING

Denote Γ the matrix with element 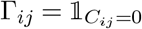 and note that the derivatives of the log-link 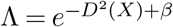 w.r.t. *D*^2^(*X*) and *β* are

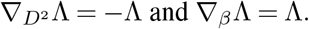

We will use these relations to derive the update steps for the optimization algorithm.

## D.1. Hurdle Poisson

Recall that the loss function has the following form

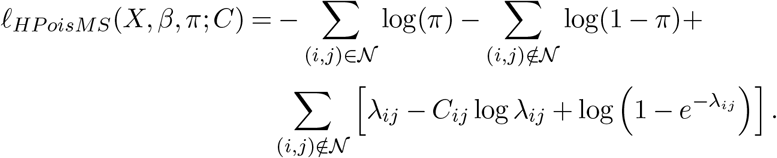

We first compute the derivatives of the loss w.r.t. *D*^2^. Denote 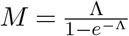, then

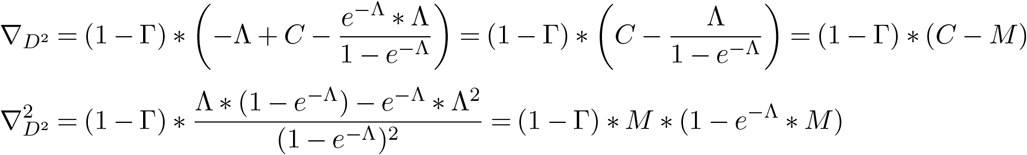

Thus we can calculate the weights and the working response for the second-order approximation and apply WPCMS to update the reconstruction *X*.

Now we deal with the nuisance parameters. First, by analogy with the distance matrix one can compute the first and second order derivatives w.r.t. *β*

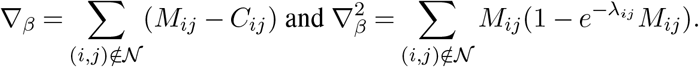

We will use these derivatives to find optimal *β* when applying Newton’s method. Next, we note that the optimal value for the Bernoulli parameter *π* can be found from the equation

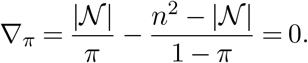

This leads us to the explicit formula 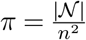. Combining all the above steps leads us to the following *HPoisMS algorithm*.

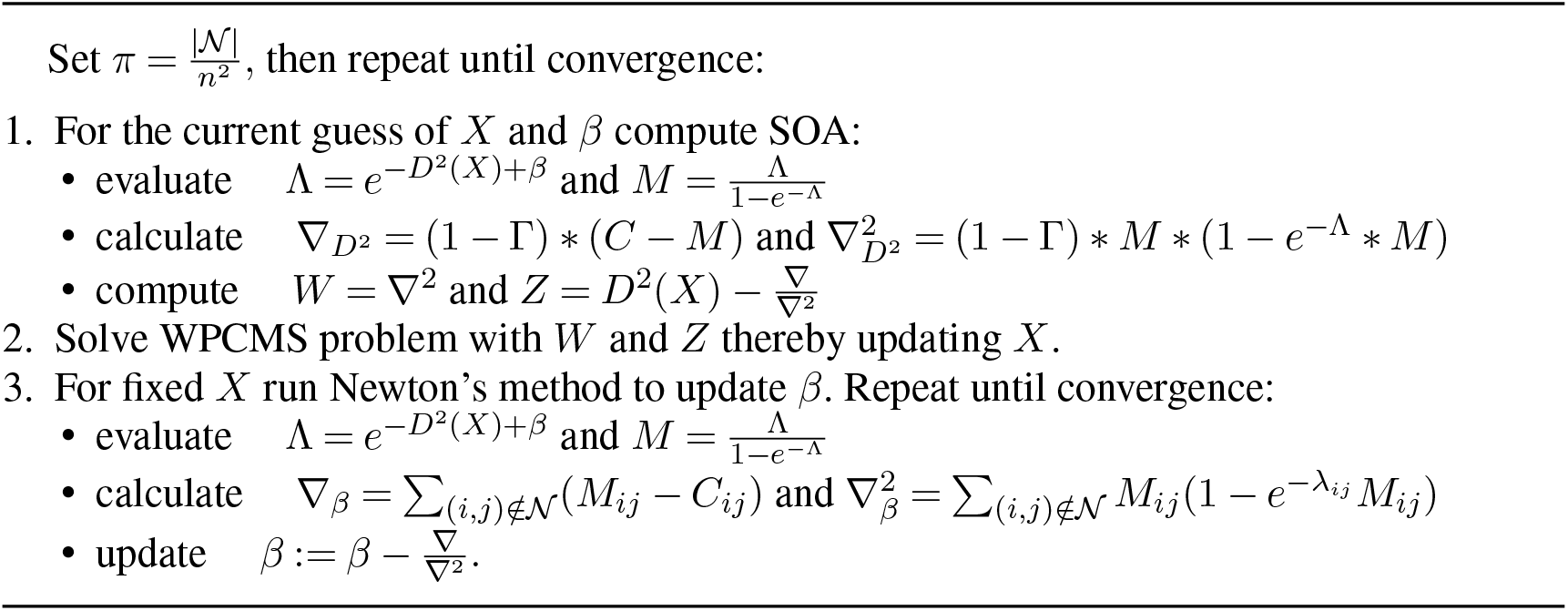
Hurdle Poisson Metric Scaling (HPoisMS)

## D.2. Zero-inflated Poisson

In this section be build an optimization algorithm for the zero-inflated model. Let 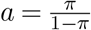 and rewrite the ZIPoisMS loss function as

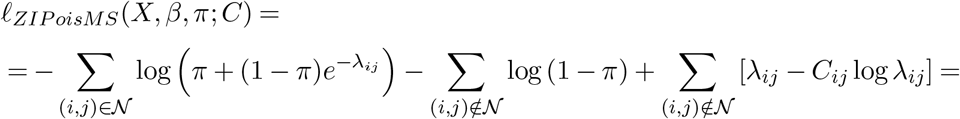

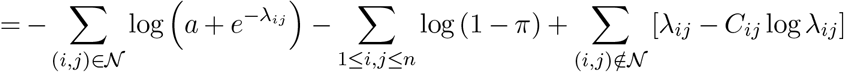

We first calculate the derivatives of the loss w.r.t. *D*^2^:

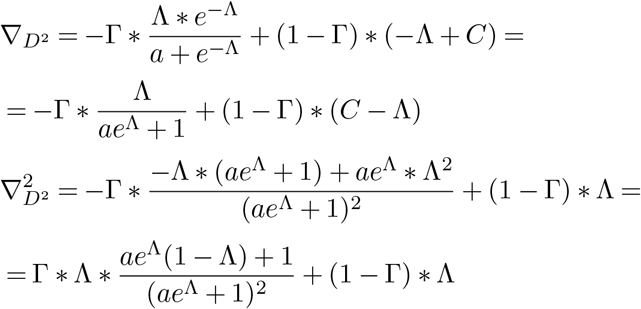

Next, by analogy, we compute the derivatives w.r.t. the intercept *β*:

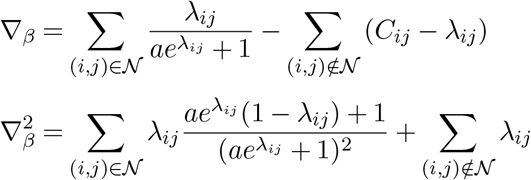

We will use these formulas to update the intercept via Newton’s method. Now, we handle the nuisance parameter *π*. Unlike the Hurdle model, there is no explicit formula for optimal *π*; therefore, we also apply Newton’s method to update it at each iteration. This requires us to compute the derivatives w.r.t. *π*

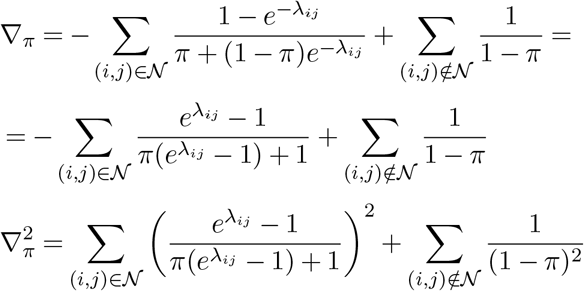

We combine all the steps in the *ZIPoisMS algorithm* stated below.

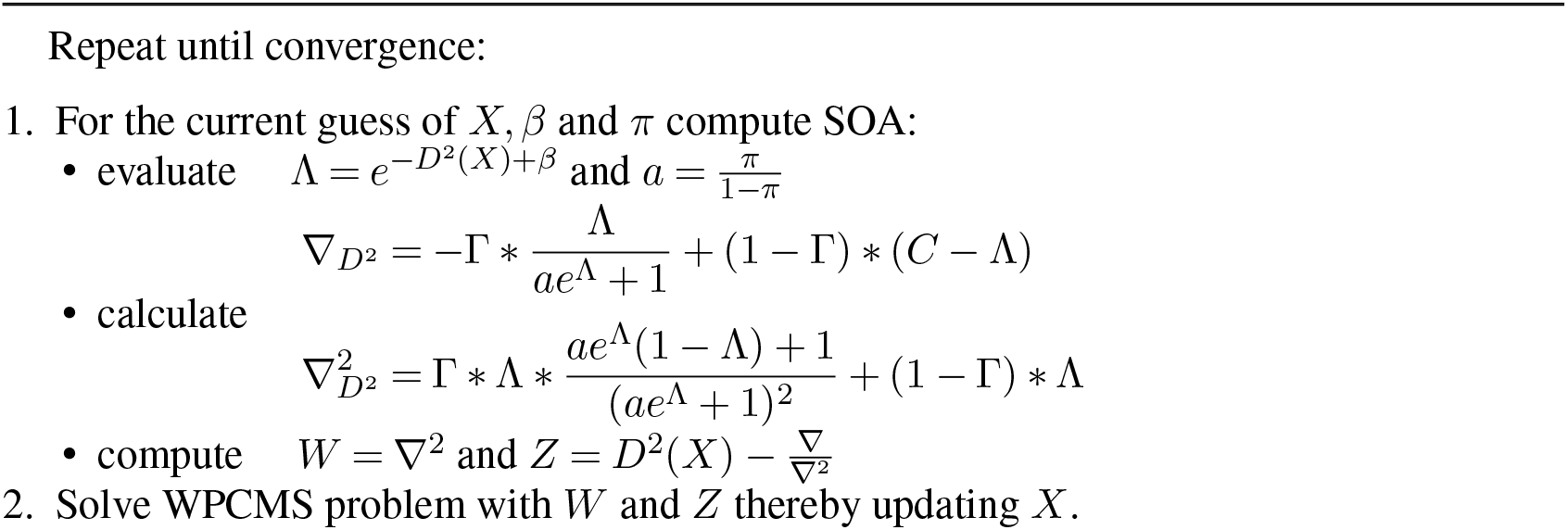
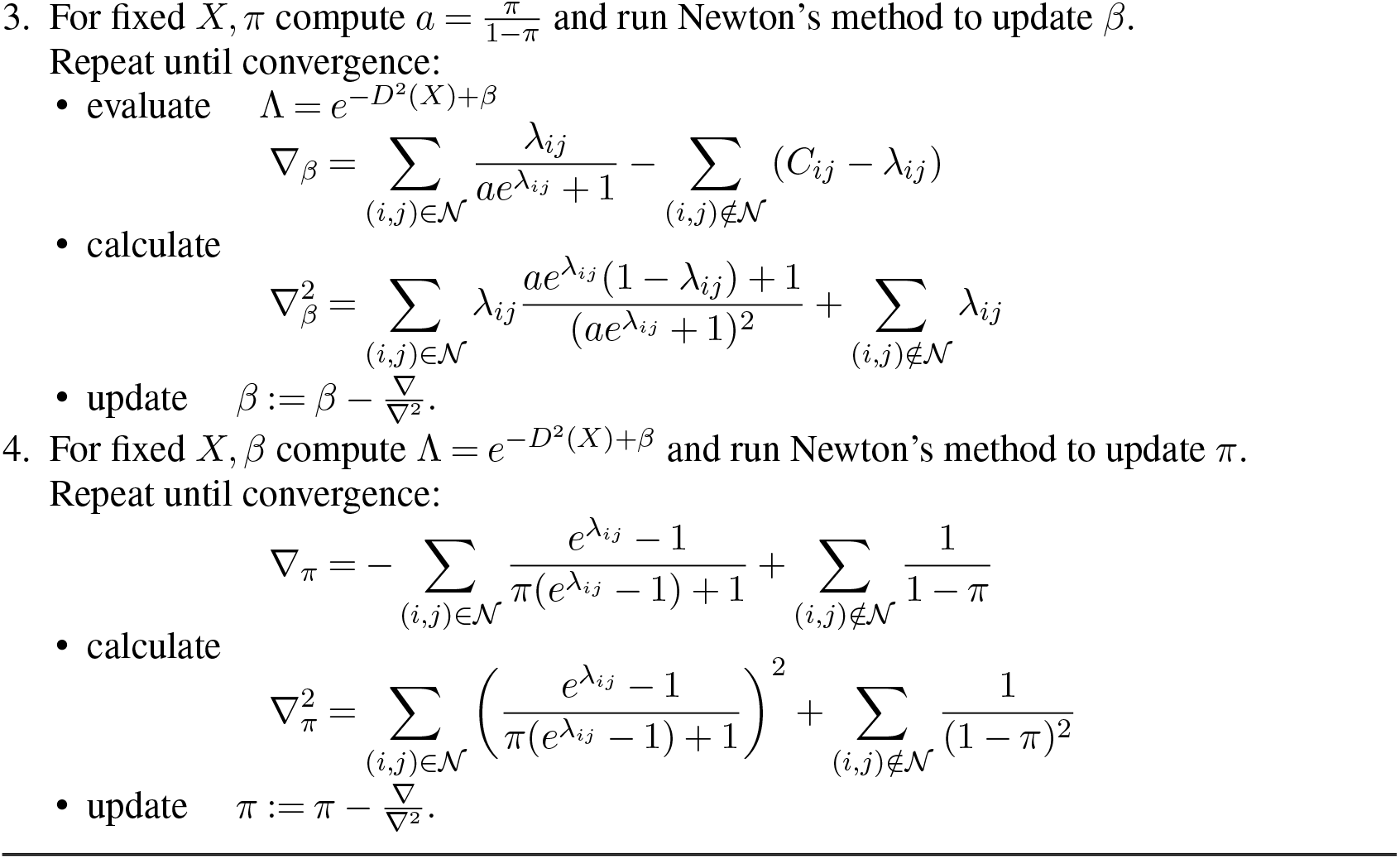
Zero-inflated Poisson Metric Scaling (ZIPoisMS)

## D.3. Negative Binomial

We extend the methodology to the negative binomial model. We start with taking the derivatives of the loss function

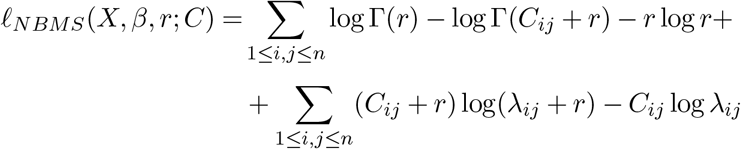

w.r.t. *D*^2^ leads us to the following equations

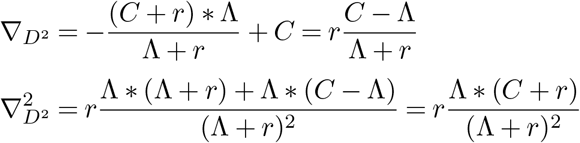

We will use these derivatives to calculate the WPCMS parameters and, subsequently, update the reconstruction *X*. Next, we calculate the derivatives w.r.t. *β*

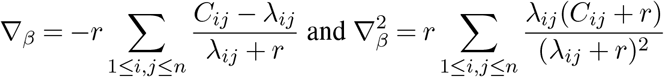

and use them to update *β* via Newton’s method. Finally, to find optimal *r* we compute the derivatives w.r.t. this nuisance parameter:

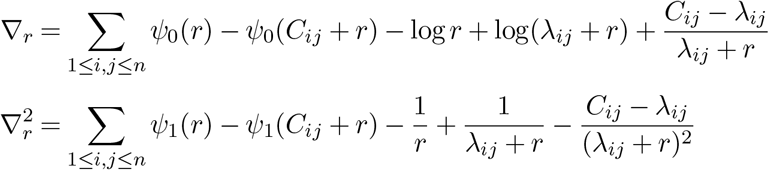

Here *ψ*_0_(.) and *ψ*_1_(.) correspond to the di- and tri-gamma function. We conclude this section with the NBMS algorithm.

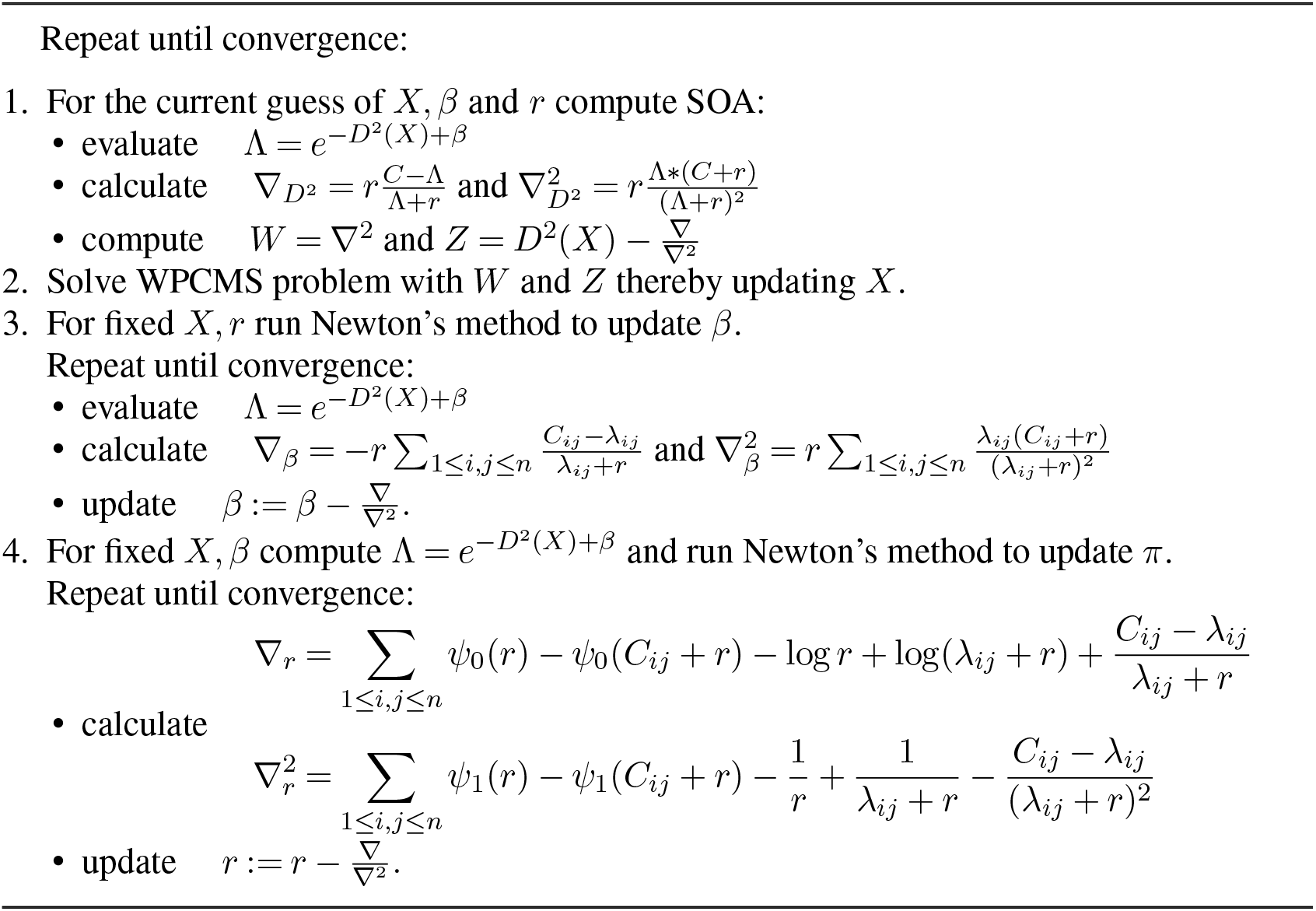
Negative Binomial Metric Scaling (NBMS)

## Notes

### Competing Interest Statement

The authors have declared no competing interest.

